# MUC5AC drives COPD exacerbation severity through amplification of virus-induced airway inflammation

**DOI:** 10.1101/706804

**Authors:** Aran Singanayagam, Joseph Footitt, Benjamin T Kasdorf, Matthias Marczynski, Michael T Cross, Lydia J Finney, Maria-Belen Trujillo Torralbo, Maria Calderazzo, Jie Zhu, Julia Aniscenko, Thomas B Clarke, Philip L Molyneaux, Nathan W Bartlett, Miriam F Moffatt, William O Cookson, Jadwiga Wedzicha, Christopher M Evans, Oliver Lieleg, Patrick Mallia, Sebastian L Johnston

## Abstract

The respiratory tract surface is protected from inhaled pathogens by a secreted layer of mucus that is rich in mucin glycoproteins. Disrupted mucus production is a cardinal feature of chronic respiratory diseases but how this alteration affect interactions between mucins and pathogens is complex and poorly understood. Here, we identify a central and unexpected role for the major airway mucin MUC5AC in pathogenesis of virus-induced exacerbations of chronic obstructive pulmonary disease (COPD). Virus induction of MUC5AC is augmented in COPD compared to healthy subjects, is enhanced in frequent exacerbators and correlates with inflammation, symptom severity and secondary bacterial infection during exacerbation. MUC5AC is functionally related to inflammation as MUC5AC-deficient (*Muc5ac*-/-) mice had attenuated rhinovirus-induced airway inflammation whilst exogenous MUC5AC glycoprotein administration augmented virus-induced inflammatory responses and bacterial load. Mechanistically, MUC5AC-augmentation of rhinovirus-induced inflammation occurred through release of extracellular adenosine triphosphate (ATP). Therapeutic suppression of virus-induced MUC5AC release using an epidermal growth factor receptor (EGFR) inhibitor ameliorated exaggerated pro-inflammatory responses in a mouse COPD exacerbation model. Collectively, these studies demonstrate previously unrecognised pro-inflammatory effects of MUC5AC during infection and thus highlight a key unforeseen role in driving COPD exacerbation severity.

## INTRODUCTION

Chronic obstructive pulmonary disease (COPD) is an inflammatory airway disorder punctuated by acute exacerbations, frequently precipitated by rhinoviruses (RVs)^1,2^. Exacerbations are the major cause of morbidity and mortality in COPD^3,4^ and a greater understanding of the underlying pathophysiological mechanisms is urgently needed to facilitate development of new therapies to reduce the impact of these debilitating episodes.

Respiratory mucosal surfaces are continuously exposed to inhaled pathogens and a protective layer of secreted mucus that is rich in mucin glycoproteins acts as a first-line of defense against infection. Alterations in mucus secretion including enhancement of the major airway mucins MUC5AC and MUC5B are a cardinal feature of inflammatory airway COPD^5–7^. The complexity of interactions between mucins and pathogens and how the perturbations in mucin expression that occur in COPD affect infection susceptibility and subsequent exacerbation pathogenesis are poorly understood and it is not known whether inhibition of mucus production would be beneficial or harmful in this context.

Animal studies have revealed critical roles for MUC5AC and MUC5B in anti-viral and anti-bacterial host-defense ^8,9^ suggesting that use of anti-secretory therapies may compromise a protective role for mucins during exacerbations. Conversely, mucin levels are higher in COPD patients who experience frequent exacerbations ^7^ raising speculation for a possible functional role in exacerbation pathogenesis. Additionally, accumulation of small airway mucus-containing inflammatory exudates has been shown to be associated with early death in patients with COPD treated by lung volume reduction surgery ^10^ and airway MUC5AC mediates airway hyperresponsiveness ^11^ and is over-expressed in cases of fatal asthma ^12^, suggesting that mucins could be a marker or cause of adverse outcome. These studies would support the opposing view that targeted suppression of mucins during exacerbations might provide desirable improvement in outcomes.

Here, using human studies in combination with functional experiments in mouse models, we identify a central and unforeseen role for MUC5AC in amplifying virus-induced airway inflammation and driving subsequent exacerbation severity in COPD. We identify a mechanism for augmentation of virus-induced inflammation by MUC5AC through release of extracellular adenosine triphosphate (ATP) and show that therapeutic suppression of virus-induced MUC5AC release using an epidermal growth factor receptor (EGFR) inhibitor ameliorates exaggerated pro-inflammatory responses in a mouse COPD exacerbation model. Our studies are the first to implicate MUC5AC as a key functional driver of COPD exacerbation pathogenesis and indicate that selective therapeutic inhibition of this mucin could lead to improved clinical outcomes.

## RESULTS

### MUC5AC is induced during naturally-occurring COPD exacerbations and enhanced in frequent exacerbators

We initially evaluated the expression of the major mucin glycoproteins MUC5AC and MUC5B in sputum samples taken from COPD patients during virus-associated naturally-occurring exacerbations (**Fig.1a**) in our community-based cohort study. A total of 27 exacerbations were reported, of which 18 exacerbations (67%) were associated with positive virus detection by PCR of sputum samples (rhinovirus n=11, coronavirus n=4, RSV n=2, influenza n=1) as previously reported.^13^ These 18 virus-positive exacerbation samples were used for analyses (see **Table 1** for the clinical characteristics of the subjects reporting these 18 exacerbations). Significant increases from baseline were observed for sputum supernatant MUC5AC protein concentrations at exacerbation presentation and at 2 weeks during exacerbation with no significant increase observed for MUC5B protein (**Fig.1b**). During exacerbation, peak concentrations of MUC5AC (i.e. maximal concentrations detected during exacerbation) protein were increased compared to baseline with no significant increase observed for MUC5B (**Fig.1c**). Frequent exacerbators (patients with a history of ≥2 exacerbations in the preceding year) had increased sputum MUC5AC concentrations at exacerbation presentation and at 2 weeks during exacerbation compared with infrequent exacerbators (0 or 1 exacerbations in the preceding year), with no significant differences observed for MUC5B (**Fig. 1d**).

**Figure 1:**
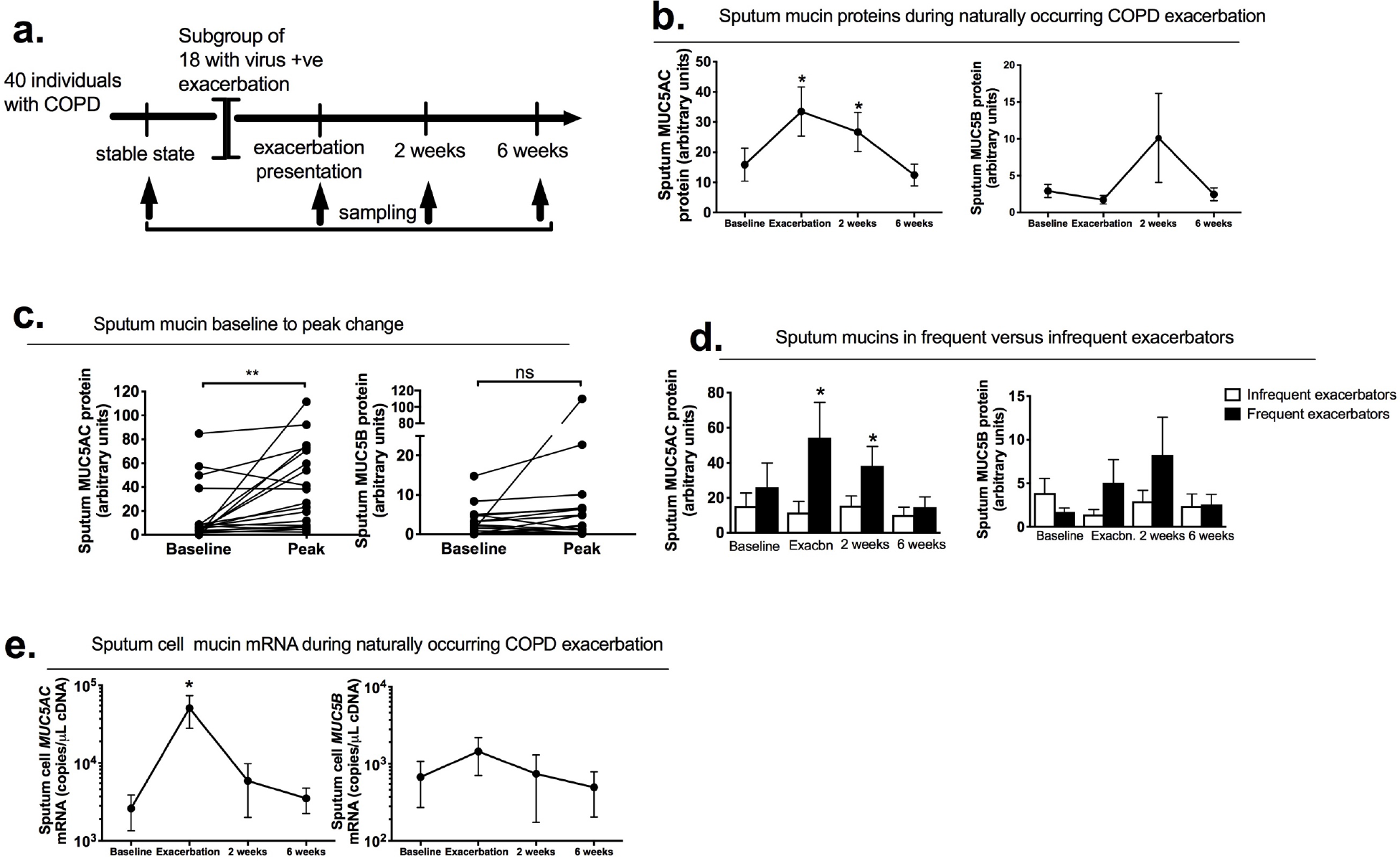
Airway mucin expression during naturally occurring COPD exacerbation. A cohort of 40 patients with COPD was monitored prospectively. Sputum samples were taken during stable state (baseline), at presentation with exacerbation associated with positive virus detection, 2 and 6 weeks following exacerbation presentation. (b) Sputum MUC5AC and MUC5B proteins. (c) baseline and peak (i.e., the maximal concentrations of mucin detected during the infection for each individual) levels of MUC5AC and MUC5B protein during naturally occurring exacerbation. (d) Sputum MUC5AC and MUC5B concentrations in frequent versus infrequent COPD exacerbators. (e) Sputum cell *MUC5AC* and *MUC5B* mRNA expression. Data analysed by Mann-Witney U test or Wilcoxon matched-pairs signed rank test. \**P*<0.05, \*\**P*<0.01.

**Table 1:** Clinical characteristics of COPD subjects with naturally occurring exacerbations associated with positive virus detection. GOLD = Global Initiative for Obstructive Lung Disease.

|  |  |
| --- | --- |
| Age (median (IQR)) (years) | 68 (63-74) |
| Male:Female sex | 11:7 |
| GOLD stage<br>I/II/III/IV | 3/11/3/1 |
| Current smoker | 5 (27.8%) |
| Ischaemic heart disease | 1 (5.6%) |
| Diabetes mellitus | 2 (11.1%) |
| Cerebrovascular disease | 1 (5.6%) |
| Inhaled corticosteroid use | 7 (38.9%) |
| Beta agonist inhaler use | 5 (27.8%) |
| Anti-muscarinic inhaler use | 4 (22.2%) |
| Frequent exacerbator ($\geq 2$ in<br>preceding year) | 7 (38.9%) |
| Prednisolone initiated for<br>exacerbation | 3 (16.7%) |
| Antibiotics initiated for exacerbation | 8 (44.4%) |

Given that ELISA-based quantification of sputum mucins may be influenced by proteases present in this sample type ^14^, we further confirmed our observations using quantitative PCR to measure mucin expression in RNA extracted from sputum cells from the same subjects. In keeping with our findings for mucin proteins, we also observed that sputum *MUC5AC* mRNA was significantly increased at exacerbation onset with no increase observed for *MUC5B* mRNA (**Fig 1e**), providing additional evidence that MUC5AC is the major induced mucin during naturally-occurring COPD exacerbations.

### COPD is associated with augmented MUC5AC responses to experimental rhinovirus challenge

We next utilised our previous human RV challenge studies^1,15^ to more accurately define the temporal dynamics of mucin expression during RV-induced COPD exacerbations (**Fig.2a** and **Supplementary Fig.1a**)(Clinical characteristics of these subjects are shown in Table 2). A significant increase in sputum MUC5AC from baseline was observed in RV-infected COPD subjects at day 3 post-infection with no increase observed in healthy control subjects (**Fig.2b**). MUC5AC was increased in patients with COPD versus healthy controls on days 3, 9 and 12 post-infection (**Fig.2b**). There was no significant change in MUC5B from baseline at any time point in patients with COPD or healthy controls and no difference in MUC5B was observed between these two groups during infection (**Supplementary Fig.1b**). MUC5AC peak concentrations were increased in patients with COPD compared to baseline with no significant change observed in healthy individuals and peak MUC5AC concentrations were higher in COPD compared to healthy subjects (**Fig.2c**). MUC5B peak concentrations were increased in both groups of volunteers when compared to baseline with no significant difference observed between the two groups (**Supplementary Fig.1c**).

**Figure 2:**
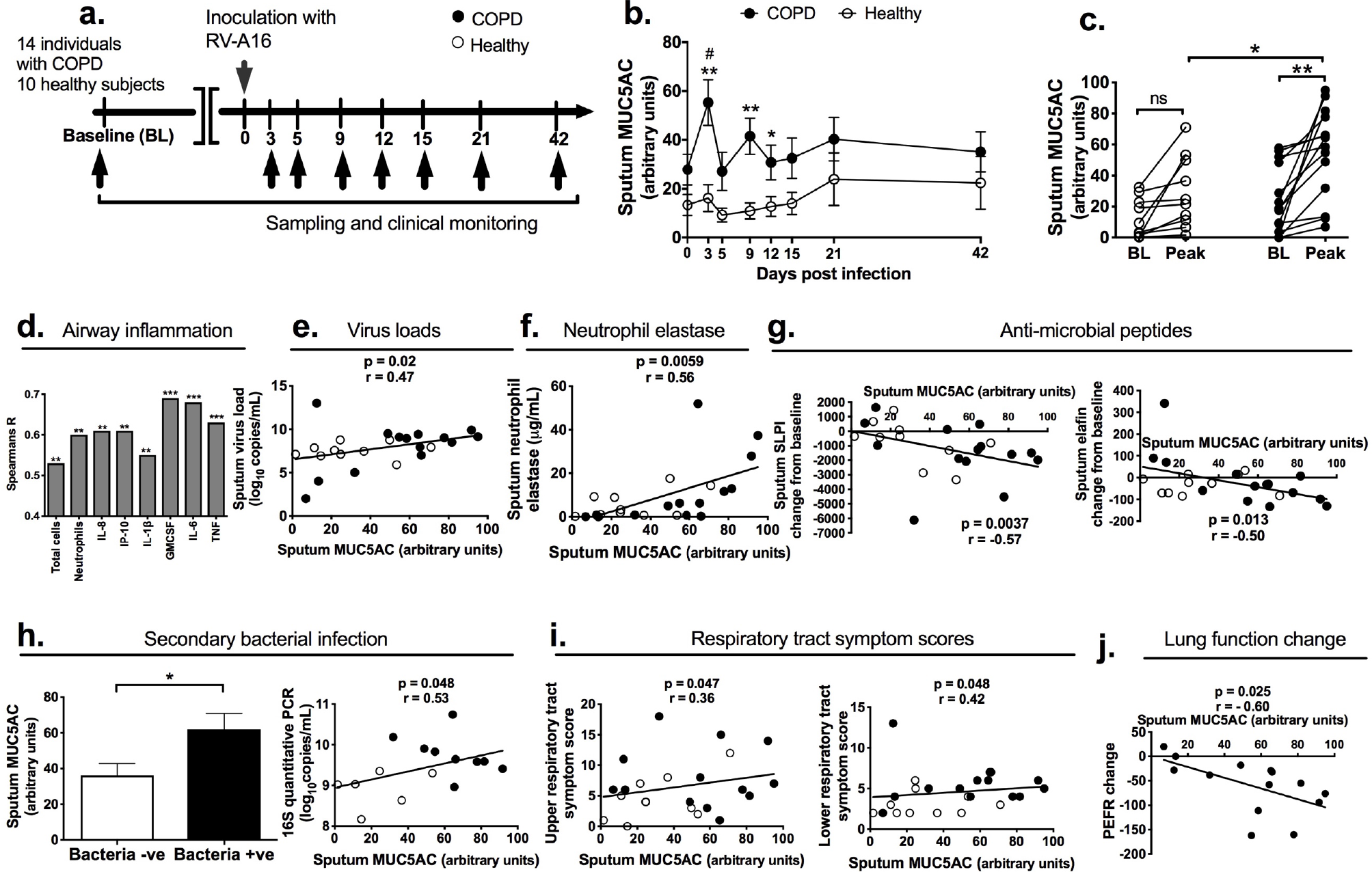
Airway MUC5AC expression and correlation with inflammatory parameters during experimental rhinovirus infection in COPD and healthy subjects. (a) Experimental outline. 14 subjects with COPD and 10 healthy control volunteers underwent sampling and analysis at baseline (̃ 14 d before RV-A16 inoculation) and at the indicated timepoints after RV-A16 infection. (b) Sputum MUC5AC concentrations in COPD and healthy subjects at baseline, and after RV-A16 infection, measured over time (c) Comparison of baseline and peak (i.e. the maximal concentration of MUC5AC detected during the infection for each individual) levels of MUC5AC and between subjects with COPD and healthy subjects. (d) Correlation of peak sputum MUC5AC concentrations with peak numbers of inflammatory cells and concentrations of cytokines in sputum. (e) Correlation between peak sputum MUC5AC and peak sputum virus loads. (f) Correlation between peak sputum MUC5AC and peak neutrophil elastase. (g) Correlation of peak MUC5AC with peak change from baseline concentrations of the indicated anti-microbial peptides in sputum. (h) Left panel: sputum MUC5AC concentrations in COPD patients with positive and negative sputum cultures for bacteria during experimental RV infection; Right panel: Correlation of peak sputum MUC5AC with bacterial loads assessed by 16S quantitative PCR. (i) Correlation of peak MUC5AC with peak respiratory tract symptom scores. (j) Correlation of peak sputum MUC5AC with peak expiratory flow rate (PEF) change from baseline during exacerbation. In (b)\**P*<0.05 \*\**P*<0.01 (comparison COPD versus healthy) *#P*<0.05 (comparison of day 3 versus baseline in COPD patients). In (c) individual datapoints shown, analysed by Wilcoxon matched-pairs signed rank test for baseline vs peak and Mann Witney U test for comparison of peak values between COPD and healthy subjects. \**P*<0.05 \*\**P*<0.01. In (h), left panel, data analysed by Mann Witney U test, \**P*<0.05. In all other panels, correlation analysis used was nonparametric (Spearman’s correlation) performed on healthy volunteers and individuals with COPD pooled into a single group. In (h), right panel, 16S qPCR not measured in all patients due to lack of sample availability so data only shown where measured.

**Table 2:**
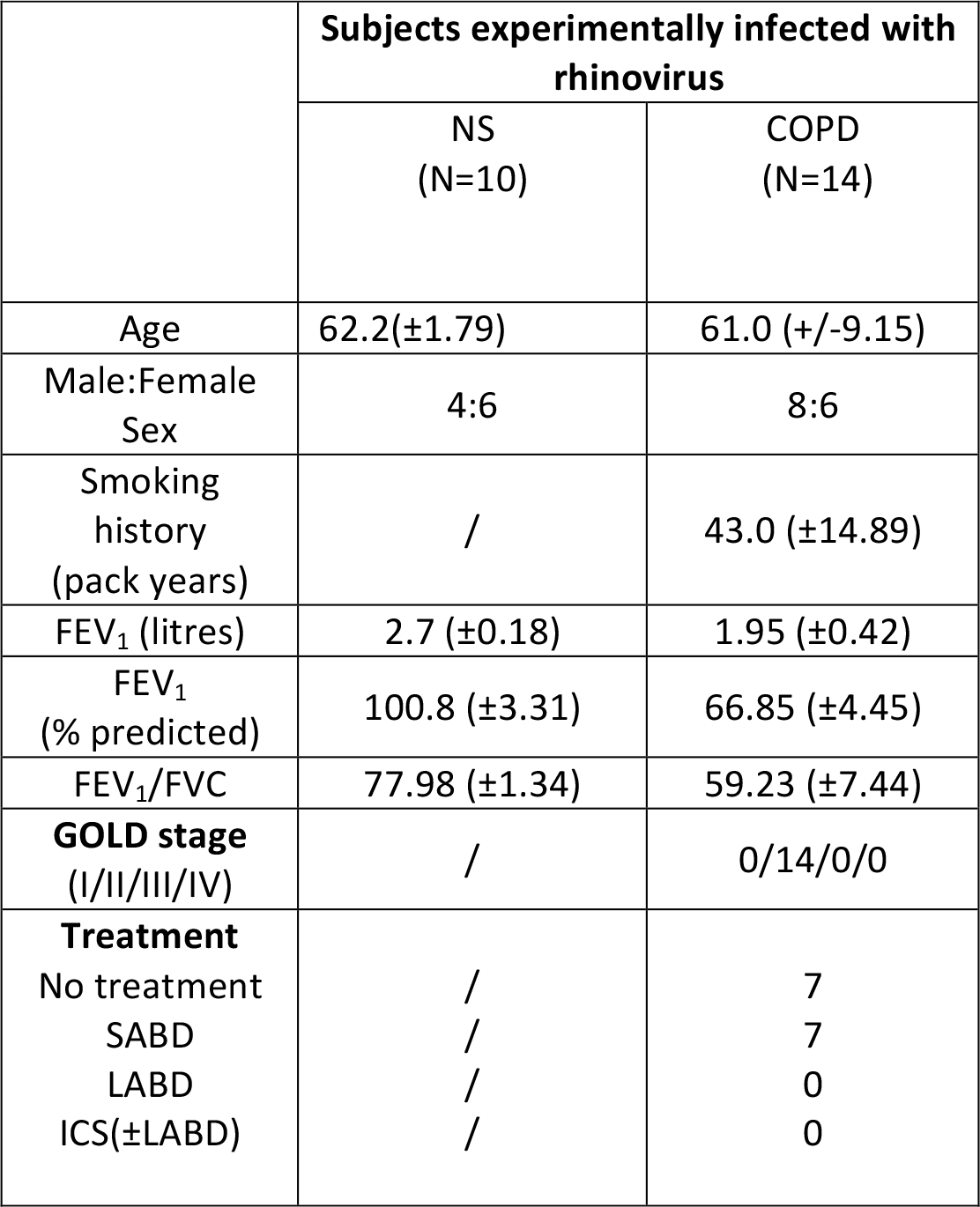
Clinical characteristics of study subjects. NS - non-smokers, COPD - Chronic Obstructive Pulmonary Disease, FEV_1_ - Forced Expiratory Volume in 1 second, FVC - Forced Vital Capacity, GOLD - Global Initiative for Obstructive Lung Disease, SABD - short-acting bronchodilators, LABD - long-acting bronchodilators, ICS - inhaled corticosteroids. Data shown is number of subjects or mean (SD).

### RV-induced MUC5AC is related to COPD exacerbation severity

Having characterised the expression of mucins during exacerbation, and noting that MUC5AC induction occurs at day 3, a time-point that precedes peak airway inflammation, which occurs between days 9 and 15 post-infection^1,15^, we next sought to assess relationships between MUC5AC responses and inflammatory responses to infection in our human experimental challenge model. We observed highly significant positive correlations between sputum MUC5AC and cellular airway inflammation (total cell counts and neutrophil counts, **Fig.2d**) and between MUC5AC and concentrations of soluble mediators of inflammation including CXCL8/IL-8, CXCL10/IP-10, IL-1β, GM-CSF IL-6 and TNF (**Fig.2d**). Since virus replication acts as a driver for airway inflammation^1,15^, we also examined relationships between sputum mucins and virus load and found a positive correlation between sputum MUC5AC and sputum virus load (**Fig.2e**).

We have previously reported that rhinovirus infection induces secondary bacterial infection in COPD through neutrophil elastase cleaving the anti-microbial peptides (AMPs) secretory leucocyte proteinase inhibitor (SLPI) and elafin to promote bacterial growth^16^. We found that sputum MUC5AC correlated positively with sputum neutrophil elastase concentrations during infection (**Fig.2f**) and negatively with the change from baseline in sputum SLPI and elafin concentrations during infection (**Fig.2g**), consistent with greater MUC5AC levels during infection being associated with greater induction of neutrophil elastase, and consequent increased degradation of these AMPs. COPD patients with positive bacterial cultures during RV infection also had higher concentrations of sputum MUC5AC during infection and sputum MUC5AC concentrations correlated with bacterial loads determined by 16S qPCR (**Fig.2h**). To confirm these latter findings observed in the rhinovirus challenge model, we performed similar analyses in the naturally occurring COPD exacerbation study and found that sputum MUC5AC concentrations during virus-induced exacerbations also correlated with bacterial loads in the 2 week sample during exacerbation, as the peak of secondary bacterial infections occurs around 2 weeks after virus infection^16^ (**Supplementary Fig.2a**). Although not induced during infection at any time point assessed, similar analyses were carried out for MUC5B. However, in contrast to MUC5AC, MUC5B did not significantly correlate with any inflammatory marker except GM-CSF (**Supplementary Fig.1d**). Similarly, there were no significant correlations of MUC5B with virus load (**Supplementary Fig.1e**), and changes from baseline in antimicrobial peptide concentrations (**Supplementary Fig. 1f**) or bacterial loads during experimental (**Supplementary Fig.1g**) or naturally-occurring (**Supplementary Fig.2b**) COPD exacerbation.

Pro-inflammatory cytokines and cellular airway inflammation are believed to contribute to duration and severity of RV-induced asthma exacerbations^17,18^ and secondary bacterial infection is also associated with greater exacerbation severity in RV-induced COPD exacerbations^16^. Having observed associations between concentrations of MUC5AC and airway inflammation, virus load and secondary bacterial infections, we next determined whether airway mucins during exacerbations were related to clinical outcomes during exacerbation. We observed that sputum MUC5AC concentrations correlated with both upper and lower respiratory tract symptom scores (**Fig.2i**) with no significant correlations observed for MUC5B (**Supplementary Fig.1h**). Since chronic mucus hypersecretion is associated with long term FEV_1_ decline^19^, we also hypothesized that the amplified induction of MUC5AC during RV infection would be related to acute lung function changes during COPD exacerbation. In experimental RV-induced COPD exacerbations, sputum MUC5AC concentrations correlated negatively with the maximal fall from baseline in peak expiratory flow during exacerbation (**Fig. 2j**) with no such correlation observed for MUC5B (**Supplementary Fig.1i**).

To confirm these findings, we additionally measured MUC5AC in an alternative sample type, bronchoalveolar lavage (BAL), as subjects in the experimental infection study also underwent bronchoscopy at baseline and day 7 after RV infection. BAL MUC5AC protein was similarly increased in COPD subjects versus healthy non-smokers with no difference observed for MUC5B (**Supplementary Fig 3**).

Combined, these observations confirmed that MUC5AC is the major induced mucin during human virus-induced COPD exacerbations and that concentrations of MUC5AC correlate with inflammation, virus load, secondary bacterial infections and clinical measures of exacerbation severity.

### *Muc5ac*−/− mice have attenuated inflammatory responses to RV infection

We next sought to further understand the functional importance of MUC5AC during RV infection using mouse models in which cause and effect relationships can readily be assessed. Mice with gene-targeted deletion of *Muc5ac* (*Muc5ac*-/-) (**Fig.3a**) had attenuated cellular airway inflammation (BAL total cell counts and neutrophil numbers; **Fig.3b**) and reduced concentrations of the neutrophil chemokines CXCL1/KC and CXCL2/MIP-2 (**Fig. 3c**) following RV infection compared to wild-type controls. BAL concentrations of the pro-inflammatory cytokines IL-1β, IL-6 and TNF were also reduced in RV-infected *Muc5ac*-/- mice compared to wild-type controls (**Fig. 3d**). The pro-inflammatory effects of MUC5AC were not mediated via interference with antiviral immune responses or virus control as *Muc5ac*-/- mice and wild type controls had similar levels of IFN-α and IFN-λ2/3 protein in BAL and no difference in lung virus loads (**Supplementary Fig 4a-d**). These data indicated that MUC5AC is functionally related to airway inflammatory responses during RV infection.

**Figure 3:**
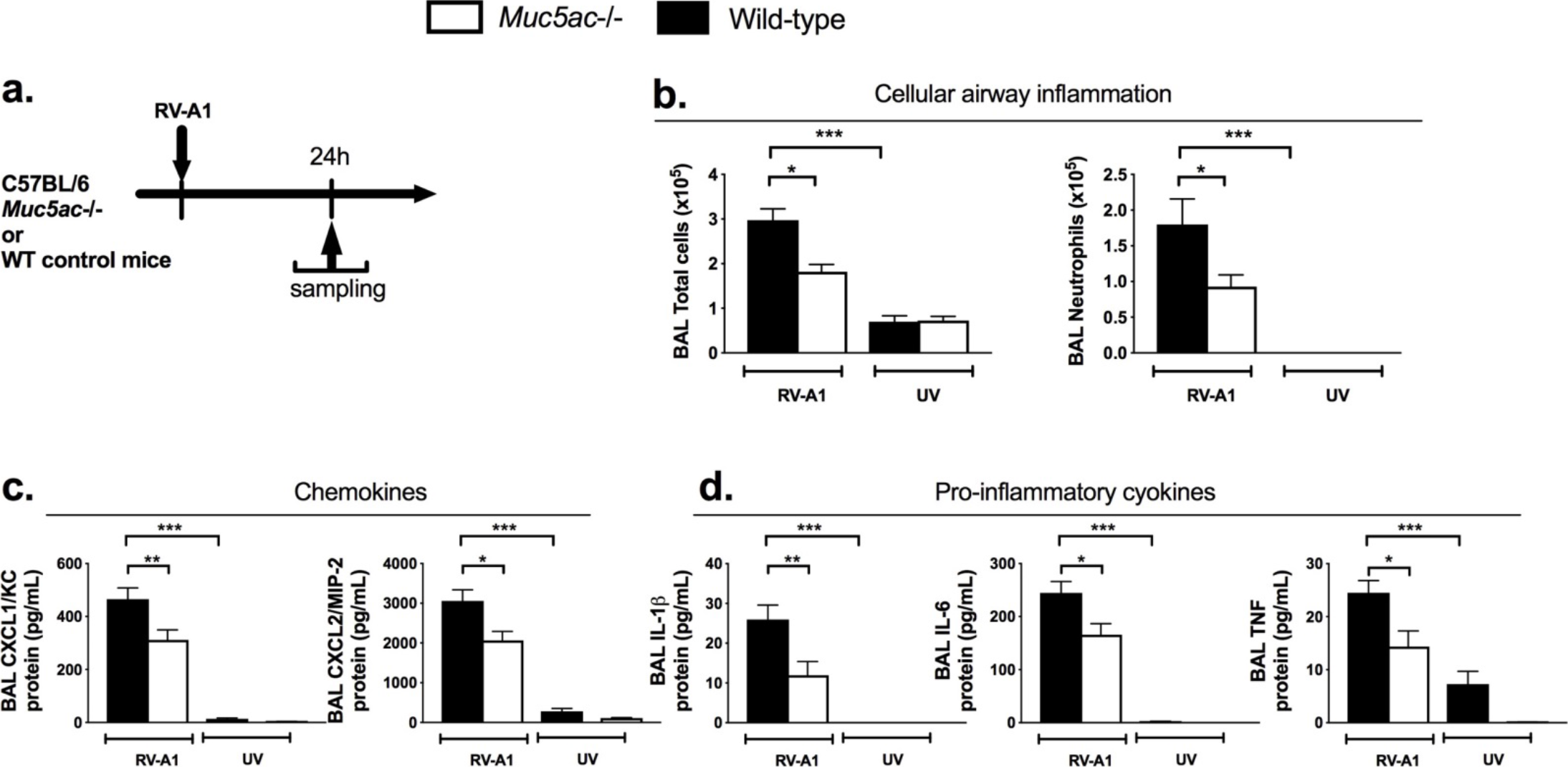
*Muc5ac*-/- mice have attenuated inflammatory responses to RV infection. (a) Experimental outline. *Muc5ac*-/- or wild-type control C57BL/6 mice were intranasally infected with RV-A1 or UV inactivated RV-A1. BAL total cells and neutrophils at day 1 post-infection. (c) BAL concentration of the indicated chemokines at day 1 post-infection. (d) BAL concentration of the indicated pro-inflammatory cytokines at day 1 post-infection. All data expressed as mean (+/−S.E.M). Data analysed by one-way ANOVA with Bonferroni’s post-test. \**P*<0.05; \*\**P*<0.01; \*\*\**P*<0.001.

### Exogenous MUC5AC protein augments RV-induced airway inflammation and increases bacterial loads in mice

To further confirm a functional role for MUC5AC in RV-induced inflammation and exacerbation severity, we evaluated the *in vivo* effects of exogenous MUC5AC glycoprotein in the mouse model of RV infection (**Fig.4a**). Exogenous MUC5AC administration alone had no effect on BAL inflammatory cell numbers (**Fig.4b**) or pro-inflammatory chemokine or cytokine concentrations (**Fig.4c&d**) but MUC5AC administration (0.5mg/mL) enhanced RV-induced airway inflammation including increased total cell and neutrophil numbers (**Fig. 4b**), and increased BAL concentrations of the pro-inflammatory chemokines CXCL1/KC, CXCL2/MIP-2 and CCL5/RANTES and cytokines IL-1β, IL-6 and TNF (**Fig.4c&d**). Conversely, administration of MUC5B protein or a control polymer solution of agarose/dextran at the same concentration (0.5mg/mL) had no effect on inflammation, either alone or when administered in combination with RV infection in mice. (**Supplementary Fig 5**)

**Figure 4:**
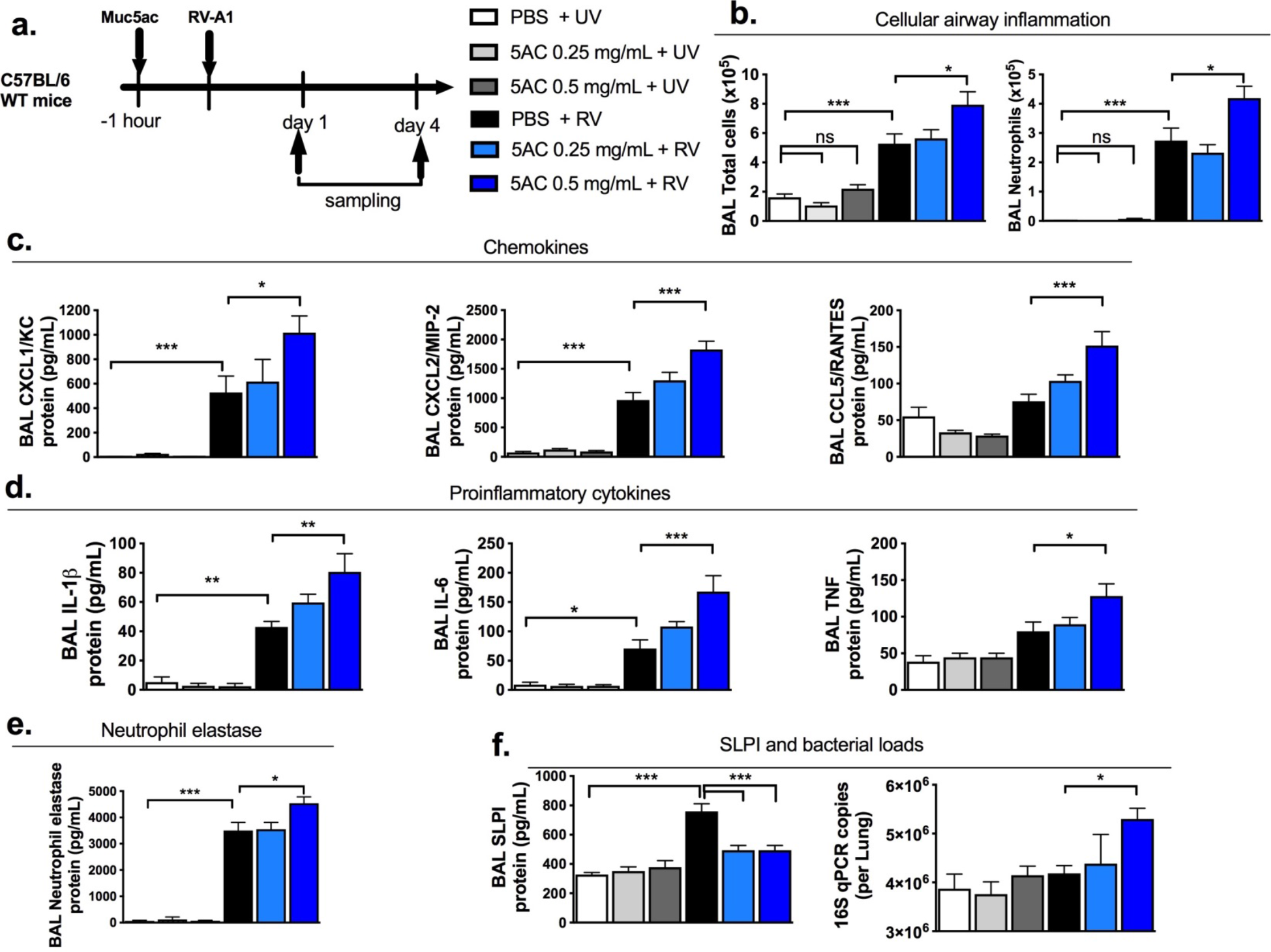
Exogenous MUC5AC augments airway inflammation and bacterial loads in rhinovirus-infected mice. (a) Experimental outline. C57BL/6 mice were treated intranasally with purified MUC5AC protein and additionally intranasally infected with RV-A1 or UV inactivated RV-A1. (b) BAL total cells and neutrophils at day 1 post-infection. (c) BAL concentrations of the indicated chemokines at day 1 post-infection. (d) BAL concentration of the indicated pro-inflammatory cytokines at day 1 post-infection. (e) BAL concentrations of neutrophil elastase at day 1 post-infection. (f) BAL concentrations of secretory leucocyte proteinase inhibitor (SLPI) and lung 16S bacterial loads at day 4 post-infection. All data expressed as mean (+/−S.E.M). Data analysed by one-way ANOVA with Bonferroni’s post-test. \**P*<0.05; \*\**P*<0.01; \*\*\**P*<0.001.

Consistent with our hypothesis that MUC5AC drives neutrophil elastase-mediated cleavage of AMPs and subsequently increases bacterial loads, we also observed that exogenous MUC5AC increased RV-induction of neutrophil elastase (**Fig.4e**), suppressed RV-induction of SLPI and augmented lung bacterial loads during virus infection (**Fig.4f**). Similar to what was observed with knockout mice, exogenous Muc5ac also had no effect on induction of IFN-α or IFN-λ2/3 nor lung virus loads in RV-infected mice (**Supplementary Fig4e-h**).

### MUC5AC exerts pro-inflammatory effects via extracellular ATP

We next sought to understand how MUC5AC amplifies inflammation during RV infection. Given our prior observations that *Muc5ac* deletion was anti-inflammatory, while augmentation with exogenous MUC5AC protein in mice had broad pro-inflammatory effects, we reasoned that this mucin could be mediating its effects via a danger signal that promotes inflammatory mediator release and cell recruitment. A previous study demonstrated that supernatants of mucopurulent material from subjects with cystic fibrosis can induce epithelial release of extracellular adenosine triphosphate (ATP)^20^, a well-recognised promoter of inflammation^21^ that is also virus-inducible^22^. We therefore hypothesised that MUC5AC exerts its pro-inflammatory effects during RV infection via induction of extracellular ATP and examined the effect of ATP neutralisation using the ATP-hydrolysing enzyme apyrase in exogenous MUC5AC-treated RV-infected mice (**Fig 5a**). Exogenous MUC5AC protein administration augmented BAL concentrations of ATP in RV-infected mice, an effect that was abrogated by instillation of apyrase (**Fig 5b**). Apyrase administration also suppressed exogenous MUC5AC-mediated increases in airway inflammation during RV infection including BAL total cells and neutrophils (**Fig. 5c**), inflammatory chemokines CXCL1/KC, CXCL2/MIP-2 and CCL5/RANTES (**Fig. 5d**) and pro-inflammatory cytokines TNF and IL-1β **(Fig. 5e**). Apyrase also prevented the exogenous MUC5AC-mediated increase in neutrophil elastase, reversed the MUC5AC-mediated suppression of SLPI and suppressed MUC5AC-mediated increased bacterial loads in RV-infected mice (**Fig 5f-g**). Apyrase-mediated reversal of pro-inflammatory effects of MUC5AC did not occur via interference with antiviral immune responses or virus control, as we also observed no effect of apyrase on RV-induction of IFN-α, IFN-λ2/3 (**Supplementary Fig.6a-c**) or lung virus loads (**Supplementary Fig.6d**).

**Figure 5:**
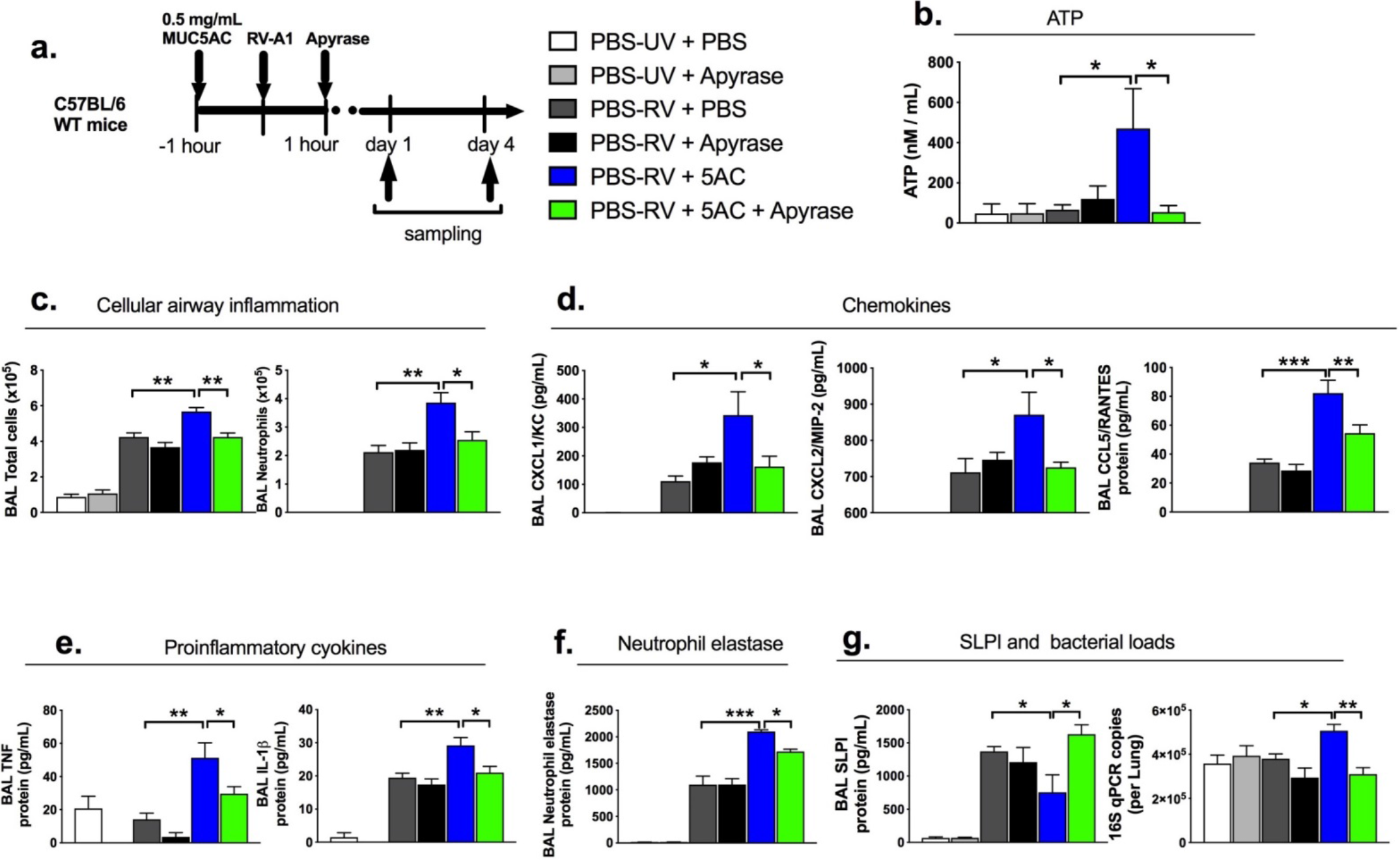
Neutralization of airway ATP levels inhibits augmentation of inflammation and bacterial loads by MUC5AC. (a) Experimental outline. C57BL/6 mice were treated intranasally with purified MUC5AC protein or PBS control, infected with RV-A1 or UV inactivated RV-A1 and additionally treated with intranasal apyrase or vehicle control. (b) BAL ATP concentrations (c) BAL total cells and neutrophils at day 1 post-infection. (d) BAL concentration of the indicated chemokines at day 1 post-infection. (e) BAL concentration of the indicated pro-inflammatory cytokines at day 1 post-infection. (f) BAL concentration of neutrophil elastase at day 1 post-infection. (g) BAL concentration of secretory leucocyte proteinase inhibitor (SLPI) and lung 16S bacterial loads at day 4 post-infection. All data expressed as mean (+/−S.E.M). Data analysed by one-way ANOVA with Bonferroni’s post-test. \**P*<0.05; \*\**P*<0.01; \*\*\**P*<0.001.

These data confirmed that MUC5AC exerts its pro-inflammatory effects during RV infection through MUC5AC-mediated induction of release of extracellular ATP.

### Therapeutic inhibition of MUC5AC induction by RV protects against RV-induced exacerbation in a mouse model of COPD-like disease

Having confirmed the functional importance of MUC5AC in RV-induced airway inflammation, we next hypothesised that inhibiting RV-induction of this glycoprotein would have beneficial effects during RV-induced COPD exacerbations. Rhinoviruses induce MUC5AC expression through EGFR signalling^23^. We therefore assessed the effect of EGFR inhibition in a mouse model of elastase-induced emphysema combined with RV-infection in which many features of human COPD exacerbation including mucus hypersecretion are recapitulated^24^. Administration of the EGFR inhibitor AG1478 prior to RV infection in elastase treated mice (**Fig.6a**) suppressed RV-induction of lung *Muc5ac* mRNA, BAL MUC5AC protein (**Fig.6b**) and suppressed airway epithelial cell mucus staining in lung sections (**Fig. 6c**, control groups are shown in **Supplementary Fig. 7**). AG1478 had no effect on BAL MUC5B protein concentrations (**Supplementary Fig.8a&b**). AG1478 administration in elastase-treated mice also suppressed RV-induction of BAL cellular airway inflammation (**Fig.6d**), as well as chemokines CXCL1/KC, CXCL2/MIP-2 and CCL5/RANTES (**Fig.6e**), pro-inflammatory cytokines IL-1β, IL-6, TNF and GM-CSF (**Fig.6f**) and neutrophil elastase (**Fig.6g**). AG1478 treatment in elastase-treated RV-infected mice also enhanced BAL SLPI levels and reduced pulmonary bacterial loads (**Fig.6h**). Finally, elastase and AG1478-treated RV-infected mice had reduced AHR to methacholine challenge compared to elastase and vehicle treated, RV infected controls (**Fig.6i**)

**Figure 6:**
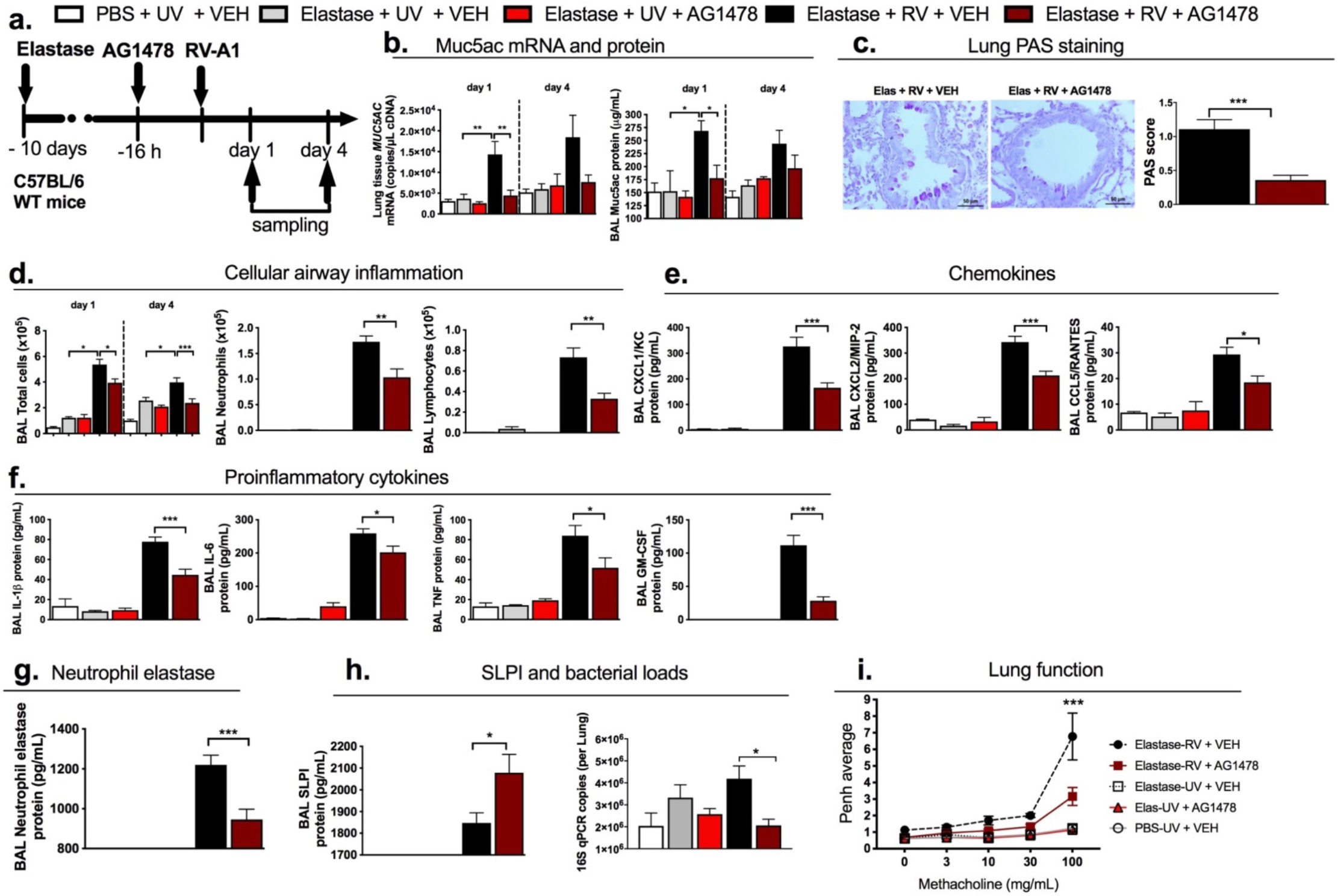
EGFR inhibition inhibits MUC5AC expression and attenuates airway inflammation, bacterial loads and airway hyper-reactivity in a mouse model of rhinovirus-induced COPD exacerbation. (a) Experimental outline. C57BL/6 mice were treated intranasally with elastase or PBS control and additionally treated intraperitoneally with 50mg/kg of EGFR inhibitor AG1478, prior to challenge with rhinovirus (RV)-A1 or UV-inactivated RV-A1 (UV). (b) Lung *Muc5ac* mRNA expression and BAL MUC5AC protein concentration. (c) Periodic-acid Schiff stained lung sections at day 4 post-infection. Left: Representative images from mice treated with elastase+RV+vehicle and elastase+RV+AG1478. Scale bars: 50μM. Magnification x400. Right: scoring for PAS-positive mucus-producing cells. (d) BAL total cells, neutrophils at day 1 and lymphocytes at day 4 post-infection. (e) BAL concentrations of the indicated chemokines at day 1 post-infection (f) BAL concentrations of the indicated pro-inflammatory cytokines at day 1 post-infection. (g) BAL concentration of neutrophil elastase at day 1. (h) BAL concentration of secretory leucocyte proteinase inhibitor (SLPI) at day 1 and lung 16S bacterial loads at day 4 post-infection. (i) Airway hyper-responsiveness to methacholine challenge at day 1 post-infection. All data expressed as mean (+/−S.E.M). Data analysed by one or two-way ANOVA with Bonferroni’s post-test. \**P*<0.05; \*\**P*<0.01; \*\*\**P*<0.001. In (i) comparison shown is for Elastase-RV + VEH versus Elastase-RV + AG1478 group.

These anti-inflammatory effects of AG1478 were not mediated via interference with antiviral immune responses or virus control, as we also observed no effect of AG1478 on RV-induction of IFN-α, IFN-λ2/3 (**Supplementary Fig.8c&d**) or lung virus loads. (**Supplementary Fig.8e**).

## DISCUSSION

The mechanisms underlying virus-induced exacerbations in COPD and the factors that drive inflammation and clinical severity during these episodes are poorly understood. Here, we provide insight into the role that the major airway mucin glycoprotein MUC5AC plays in the biology of these episodes. Using a combination of human and mouse models of rhinovirus infection in COPD, we demonstrate a novel central role of virus-induced MUC5AC in driving COPD exacerbation severity through amplification of airway inflammation, findings that open up new opportunities for targeting MUC5AC therapeutically. We indicate a mechanism for MUC5AC augmentation of RV-induced inflammation via release of extracellular ATP.

We show here, that MUC5AC is selectively induced during virus-induced COPD exacerbations and increased compared to virus-infected healthy subjects. By contrast, MUC5B was not induced above baseline and did not differ between COPD and control subjects at any timepoint, indicating that MUC5AC is the major induced mucin during exacerbations. We also found positive correlations between MUC5AC and a range of inflammatory markers. Since airway inflammation is enhanced in COPD exacerbations and believed to contribute to duration and severity of RV induced illnesses^17,18^ we also determined the relationship between MUC5AC and exacerbation severity, finding positive correlations with virus loads, symptom scores and acute lung function decline. MUC5AC is one of the major glycoprotein components of airway mucus and it is therefore unsurprising that its expression during virus infection correlates with lower respiratory symptom scores which comprise an assessment of sputum production. The correlation of MUC5AC with PEF decline can be explained by the fact that greater mucus production is likely to contribute directly to airway obstruction during exacerbation. Our findings in this acute human infection model are in keeping with clinical studies that have reported the association of symptom-defined chronic mucus hypersecretion phenotype with airflow limitation and FEV_1_ decline^19,25^ and an animal study showing that genetic deletion of *Muc5ac* reduces mucus occlusion and improves lung function in models of allergic airway inflammation^11^.

We have previously reported that experimental RV infection in patients with COPD is associated with increased frequency of secondary bacterial infection compared to healthy subjects^16,26^ via a mechanism of virus-induced neutrophil elastase-mediated cleavage of the anti-microbial peptides SLPI and elafin. The positive correlation of sputum MUC5AC with neutrophil numbers, neutrophil elastase concentrations and secondary bacterial infection and negative correlation with SLPI and elafin levels observed in the current study suggests that increased MUC5AC could be one mechanism contributing to enhanced neutrophilic airway inflammation, greater neutrophil elastase levels and thus greater risk of development of secondary bacterial infections. Siegel *et al* previously demonstrated reduced secondary pneumococcal growth following influenza infection in *Muc5ac*-/- mice^27^, an effect mediated by reduced provision of mucin-derived nutrients for bacterial growth in *Muc5ac*-/- mice, indicating that MUC5AC may be an important promoter of secondary bacterial infection via more than one mechanism. Combined with our findings in a clinically relevant human model, this highlights inhibition of MUC5AC as a potential target for reducing secondary bacterial infection in COPD.

A recent study by Kesimer *et al* showed that total mucin concentrations in sputum at stable state were higher in COPD patients who experienced frequent exacerbations^7^. Previous studies have also reported chronic mucus hypersecretion to be associated with increased exacerbation frequency^28,29^ and mortality risk related to pulmonary infection^30^. Here, we identify that frequent exacerbators also have enhanced sputum MUC5AC concentrations during exacerbation. Underlying mechanisms involved in the frequent exacerbator phenotype are poorly characterised, but our data raise speculation that these patients may have exaggerated mucin responses to infection potentially leading to enhanced airway inflammation and increased symptoms.

The human experimental challenge model facilitates sequential sampling at precisely defined time points during the course of infection and allows evaluation of the temporal relationship between variables. We observed that MUC5AC induction peaked early at day 3 in the time course of these RV-induced COPD exacerbations, preceding airway inflammation which peaks between day 9 and day 15 post-infection^1,15^. Given the strong positive correlations observed between MUC5AC and these parameters in the current study, we hypothesised that MUC5AC induced by rhinovirus might contribute directly to enhanced airway inflammation during exacerbations. As it is not possible to infer causation from measurements in our human model we carried out complementary gain and loss of function experiments in mice to directly investigate the effect of a functional role of MUC5AC during rhinovirus infection. Mice with gene targeted deletion of *Muc5ac* (*Muc5ac*-/-) had attenuated airway inflammation following RV infection, similar effects to those reported in models of ventilator-induced lung injury in the same knockout strain^31^. In our study, exogenous MUC5AC protein administration additionally augmented rhinovirus induction of pro-inflammatory responses in mice, thereby confirming that MUC5AC is functionally related to enhanced airway inflammation during virus infection, effects not observed with administration of MUC5B protein or a control polymer of agarose/dextran solution. Exogenous MUC5AC also enhanced rhinovirus-induction of neutrophil elastase, suppressed SLPI production and increased pulmonary bacterial loads, further confirming a role for MUC5AC in inhibiting anti-bacterial responses during virus infection. A previous study by Ehre *et al* evaluated responses to influenza infection in a transgenic *Muc5ac* overexpressing mouse strain and showed reduced viral titres and attenuated neutrophilic inflammation in this model^9^. This contrasts our findings using exogenous MUC5AC administration where there was no effect on virus loads and MUC5AC enhanced virus-induced inflammation. The transgenic strain employed by Ehre *et al* was associated with ~18-fold increase in airway MUC5AC levels^9^. This level of augmentation greatly exceeds the increases observed in our human analyses where a ~2-fold increase was observed in stable state levels of MUC5AC in COPD versus healthy subjects and ~2-to 3-fold increase was observed from baseline to exacerbation. The concentrations of exogenous MUC5AC protein administered in our animal experiments (0.25-0.5mg/mL) were ~ 1.5-3.5 fold higher than the baseline MUC5AC protein concentrations we measured in the mouse airway (~150μg/mL) and are therefore more likely to be physiologically relevant to human exacerbation. Further, the effect of MUC5AC in reducing virus loads at the much higher MUC5AC levels generated in the MUC5AC transgenic strain reported by Ehre *et al* may be virus-specific^9^. The authors reported that MUC5AC binds to the influenza virus receptor, α2,3-linked sialic acids in transgenic animals, consistent with a mechanism of influenza protection *in vivo* through impairment of efficient binding of virus to receptors on the bronchial epithelium leading to reduction in virus loads and consequently lesser inflammation. It is important to note that we used a non-human purified mucin protein in solution for our experiments which does not fully recapitulate the complex biophysical properties of mucin glycoprotein contained within endogenous airway mucus. We confirmed that the mucin preparation used was free of endotoxin and DNA but we cannot exclude that other mucin-bound proteins are present in the solution that may be contributing to the observed effects. Future validation of our findings using administration of human or mouse MUC5AC in similar mouse models is warranted, although extraction and purification of mucins from these sources is technically challenging and was not feasible for the current studies.

We identified a mechanism for amplification of rhinovirus-induced airway inflammation by MUC5AC through release of ATP, a danger signal that is released during infection and contributes to nucleotide receptor-dependent inflammatory responses^32^. This mechanism was suggested in a previous study that indicated that administration of sterile supernatants of mucopurulent material from subjects with cystic fibrosis can induce ATP release by human bronchial epithelial cells *in vitro*^20^. The role of ATP as a potential driver of airway inflammation in stable asthma and COPD is well recognised^33,34^. Our findings, in models of infection, that neutralisation of pulmonary ATP levels abrogates MUC5AC enhancement of inflammation and secondary bacterial infection identifies a novel mechanism through which rhinovirus-induced mucin production enhances COPD exacerbation severity.

In addition to our observation that MUC5AC directly augments rhinovirus-induced airway inflammation, a number of previous studies have also demonstrated that inflammatory mediators such as neutrophil elastase and IL-1β can directly induce MUC5AC expression in lung epithelium^35^. We therefore speculate that mucin-induced airway inflammation might trigger further production of MUC5AC leading to a vicious cycle which contributes to enhanced airway inflammation and mucus hypersecretion to drive exacerbation severity in COPD. We thus hypothesised that early targeting of MUC5AC during rhinovirus infection might beneficially interrupt this cycle and evaluated the effect of upstream inhibition of rhinovirus-induction of MUC5AC using the EGFR inhibitor AG1478 in a COPD exacerbation mouse model. AG1478 suppressed rhinovirus-induced MUC5AC levels, accompanied by reduced airway inflammation, pro-inflammatory cytokines and chemokines, neutrophil elastase and AHR, as well as enhanced SLPI production and reduced bacterial loads in rhinovirus-infected elastase-treated mice. Importantly, AG1478 had no impact on anti-viral responses, or virus control, reassuringly indicating that therapies aimed at inhibiting MUC5AC production would not be expected to adversely affect anti-viral host-defence. Jing *et al* recently reported in COPD airway epithelial cell cultures that administration of an EGFR inhibitor had no effect on RV-induction of *MUC5AC* and concluded that EGFR does not play a role in promoting virus-induced mucin expression in COPD^36^. However, in this study, the authors examined mucin expression solely at a late timepoint (15 days) post RV infection and did not evaluate effects of the treatment on mucin expression at earlier timepoints nor subsequent effects on production of inflammatory mediators. The data from our human challenge model indicates that peak RV induction of MUC5AC occurs at a much earlier timepoint (3 days) and previous studies have shown that EGFR inhibition can attenuate the early induction of MUC5AC in RV-stimulated airway epithelial cell cultures^23,37^ Our data in a mouse model of COPD-like disease supports the assertion that the early production of MUC5AC in response to RV infection occurs through EGFR and is thus amenable to therapeutic inhibition. A previous study using an inhaled EGFR inhibitor in stable COPD reported more effective EGFR inhibition was related to greater decreases in epithelial mucin stores at higher doses, but this drug was poorly tolerated^38^. Systemic EGFR antagonists are now commonly used in lung adenocarcinoma therapy^39^. Our data in mice now provides justification for human studies to evaluate the role of repurposing these agents for use in COPD exacerbations to inhibit MUC5AC production which could theoretically suppress airway inflammation, reduce secondary bacterial infection and diminish exacerbation severity.

Mucins were quantified in our study using ELISA-based assays in historically collected sputum samples specifically within a separated sputum plug aliquot, as this was the sample type available to us. Sputum plugs from severe asthmatics have previously been shown to contain more MUC5AC than MUC5B^40^ and this constituent likely contains more surface goblet cell derived mucin which may contribute to the differences observed between induction of these two mucins in our studies. However, we also observed increased RV-induced MUC5AC but not MUC5B in an alternative sample type (BAL) from COPD subjects versus healthy subjects, supporting our findings in sputum. There is also evidence to suggest that proteases within sputum samples can interfere with immunodetection of mucins^14^. Some studies have attempted to address this by using treatment with protease inhibitors shortly after sputum collection^6,41^. We have previously reported, using samples from the same study, that neutrophil elastase is increased in COPD subjects compared to healthy subjects following virus challenge^16^. Given that we observed greater induction of MUC5AC protein in COPD subjects compared to healthy subjects, despite increased levels of proteases, this suggests that the actual difference between these two groups could be even more substantial than observed in our study. Additionally, our finding that sputum *MUC5AC* but not *MUC5B* mRNA expression was increased during naturally-occurring exacerbation (mirroring what was observed at the protein level) provides an additional confirmatory approach that would not be subject to these potential confounders. Other studies have used alternative semi-quantitative antibody-independent techniques ^7,14,42^ but it is unclear whether these approaches would have sufficient sensitivity to detect subtle changes in mucin expression during infection. The primary aim of the human component of our study was to study how mucin levels change during virus-induced exacerbations and our study was unique in that we had sequential sampling from individual patients during exacerbation with consistent methods used to measure mucins in all subjects.

In conclusion, we show that MUC5AC is the major induced mucin during virus infection in COPD and that virus-induced MUC5AC plays a central role in driving airway inflammation and exacerbation severity in COPD. Future development or repurposing of therapies that specifically target this component of mucus could lead to improved clinical outcomes.

## METHODS

### Experimental rhinovirus infection studies

Samples were analysed from subjects recruited to two experimental rhinovirus infection studies that have been published previously^1,15,16^. Samples from two groups were included in this analysis: COPD subjects (GOLD stage II, not using regular inhaled therapy) and healthy non-smokers. All subjects gave informed written consent and the protocol was approved by St Mary’s NHS Trust Research Ethics Committee (study numbers 00/BA/459E and 07/H0712/138). Baseline samples of induced sputum and bronchoalveolar lavage (BAL), collected and processed as previously described^1^, were obtained when subjects had been free of COPD exacerbation/respiratory tract infection for at least 6 weeks, approximately 14 days prior to infection. Subjects were inoculated with rhinovirus-16 with samples collected on subsequent visits post-infection as described previously^1,15^. Sputum from 14 subjects with COPD and 10 non-smoking healthy control subjects had complete remaining sample availability for the full time course from baseline to day 42 and were used for the present studies.. Daily diary cards of upper and lower respiratory tract symptom scores and peak expiratory flow (PEF) measurements were taken as previously described^1^.

### The St. Mary’s Hospital Naturally-occurring COPD exacerbation cohort

A cohort of 40 COPD subjects was recruited to a longitudinal study carried out at St. Mary’s Hospital London between June 2011 and December 2013 investigating the pathogenesis of naturally-occurring exacerbations, as previously reported^43^. The subjects all had a clinical diagnosis of COPD that was confirmed with spirometry. Subjects of all grades of COPD severity were recruited and all treatments were permitted. All subjects gave informed written consent and the study protocol was approved by the East London Research Ethics Committee (Protocol number 11/LO/0229).

All subjects had an initial visit at baseline when clinically stable for clinical assessment, PEF and clinical sample collection including spontaneous or induced sputum, taken as previously described^1,44^. Subjects reported to the study team when they developed symptoms of an upper respiratory tract infection or an increase in any of the symptoms of dyspnoea, cough and sputum volume or purulence and an exacerbation was defined using the East London cohort criteria^4^. Subjects were seen within 48 hours of onset of their symptoms for collection of samples and repeat visits were scheduled for two and six weeks after the initial exacerbation visit with clinical assessment, lung function and induced sputum repeated at these time-points. Viruses were detected in sputum by PCR, as described previously^1^.

### Mouse models

Female mice (age 6-8 weeks) purchased from Charles River Laboratories on a C57BL/6 background were used for all wild-type animal studies. *Muc5ac*-/- mice were generated on a C57BL/6 background at the University of Colorado, as previously described^11^. Mice were housed in individually ventilated cages under specific pathogen-free conditions. All work was completed in accordance with UK Home office guidelines (Project licence PPL 70/7234).

For the exogenous MUC5AC protein administration model, mice were intranasally infected under light isoflurane anaesthesia with 50μL RV-A1 (5 × 10^6^ TCID_50_) or UV-inactivated RV-A1 control and additionally treated intranasally with 50μL purified porcine MUC5AC protein at 0.25 and 0.5mg/mL concentrations, 0.5mg/mL of purified MUC5B protein, 0.5mg/mL of a control polymer solution of purified agarose/dextran solution or PBS control, one hour prior to infection. Mucin proteins was prepared as previously described^45^ and confirmed to be endotoxin free using a Limulus Amebocyte Lysate (LAL) assay kit (Lonza, USA). Full details of mucin protein purification and preparation are shown in supplementary methods. In separate experiments, mice received 4 U/mL of apyrase (Sigma-Aldrich), as previously described^34^ in combination with exogenous MUC5AC and rhinovirus, one hour post-infection.

For evaluation of EGFR inhibitor in a COPD exacerbation model, as previously reported^24^, mice were treated intranasally under light isoflurane anaesthesia with 1.2 units of porcine pancreatic elastase (Merck, UK) to induce COPD-like changes. Ten days following elastase administration, as previously described^46^, mice were treated intraperitoneally with 50mg/kg of the EGFR inhibitor AG1478, 16 hours prior to intranasal infection with 50μL RV-A1 (2.5 × 10^6^ TCID_50_) or UV-inactivated RV-A1 control.

### Measurement of cytokines, mucins and ATP

Cytokine protein levels in mouse BAL, human sputum supernatants or cell supernatants from *in vitro* experiments were assayed using commercial “duoset” enzyme-linked immunosorbent assay kits (R&D Systems, Abingdon, UK). The MesoScale Discovery (MSD) platform (Maryland, USA) was used to measure inflammatory mediators IL-1β, IL-6, TNF, CXCL8/IL-8, CXCL10/IP-10 and GM-CSF in sputum supernatants according to manufacturers’ instructions as published previously^15^. Protein levels of human secretory leucocyte proteinase inhibitor (SLPI), elafin, neutrophil elastase, and all mouse BAL proteins were assayed using commercial ELISA assay kits, as previously described^16,47^.

MUC5AC and MUC5B proteins in previously stored human sputum plugs or human/mouse BAL were measured after adhesion to a 96 well plate by allowing samples to evaporate at 37°C overnight, using in-house assays as previously described^47^. For measurement of MUC5AC, the detection antibody used was biotinylated anti-MUC5AC (ThermoScientific, UK) at 400 ng/mL. For the MUC5B assay, detection antibody was mouse anti-MUC5B clone EH-Muc5ba, as previously described^48^. Bound anti-MUC5B antibody was detected with peroxidase-conjugated goat anti-mouse IgG (Sigma-Aldrich, UK). Mucins were quantified against the following standards: purified porcine MUC5AC, as previously described^45^ and recombinant MUC5B protein (Caltag Medsystems, UK).

To measure ATP levels in mouse BAL supernatants, the ATPLite assay (Perkin Elmer) was used, according to manufacturer instructions but with the cell lysis step omitted, as previously described^34^.

### RNA extraction, cDNA synthesis and quantitative PCR

For human studies, total RNA was extracted from sputum cells (RNeasy kit, Qiagen) and 2 μg was used for cDNA synthesis (Omniscript RT kit). For mouse models, total RNA was extracted from the right upper lobe of mouse lung and placed in RNA later, prior to RNA extraction and cDNA synthesis. Quantitative PCR (qPCR) was carried out using previously described specific primers and probes for human and mouse *MUC5AC and MUC5B* and rhinovirus RNA copy numbers^23,47,49^, and normalized to 18S rRNA.

### DNA extraction and bacterial 16S quantitative PCR

DNA extraction from human sputum and mouse lung was performed using the FastDNA Spin Kit for Soil (MP Biomedicals, USA), following manufacturer instructions. Bead-beating was performed for two cycles of 30 seconds at 6800 rpm (Precellys, Bertin Technologies, France). 16S bacterial loads were measured using qPCR, as previously described^50^.

### Statistical analyses

Data from human COPD studies were analysed using the Wilcoxon matched-pairs signed rank test or Mann-Witney U test. Correlations between datasets were examined using Spearman’s rank correlation coefficient. For mouse experiments, animals were studied in group sizes of 5 mice with data shown representative of at least 2 independent experiments. Data were analysed using one or two-way ANOVA with significant differences between groups assessed by Bonferroni’s multiple comparison test. All statistics were performed using GraphPad Prism 6 software. Differences were considered significant when *P*<0.05.

## Abbreviations

AMP: Anti-microbial peptide
ATP: Adenosine triphosphate
COPD: Chronic obstructive pulmonary disease
EGFR: Epidermal Growth Factor Receptor
RV: Rhinovirus

## Author contributions

**Conception and experimental design:** AS, JF, PM, SLJ

**Experimental work:** AS, JF, BTK, MM, MTC, LF, MC, MTT, JA, PLM, LG, OL.

**Writing and critical review of manuscript:** All authors

## Acknowledgements

Nil

## SUPPLEMENTARY FIGURES

**Supplementary Figure 1:**
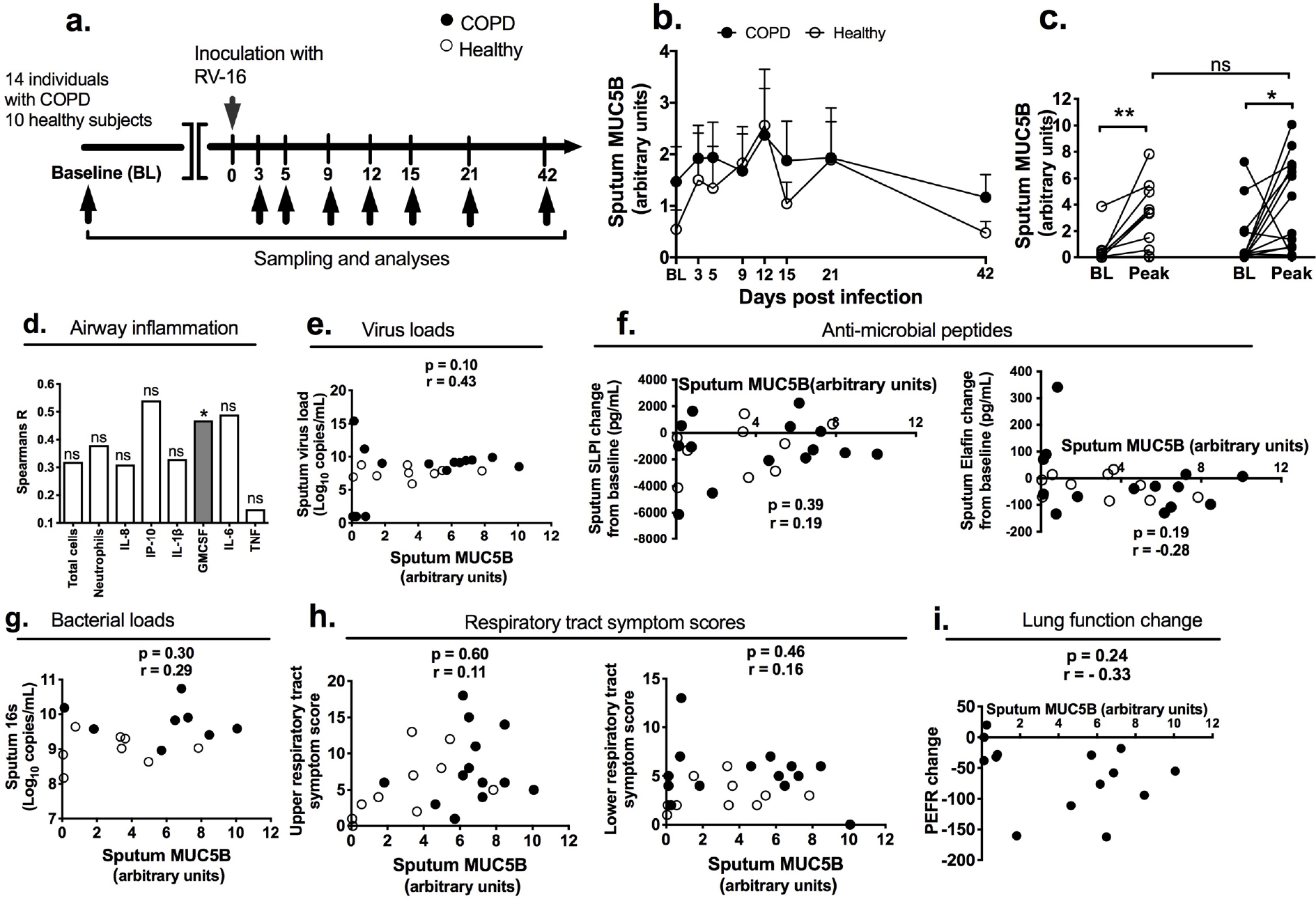
Airway MUC5B expression and correlation with inflammation and clinical parameters during experimental rhinovirus infection in COPD and healthy subjects. (a) Experimental outline. 14 subjects with COPD and 10 healthy control volunteers underwent sampling and analysis at baseline (~ 14d before RV-A16 inoculation) and at the indicated timepoints after RV-A16 infection. (b) Sputum MUC5B concentrations in COPD and healthy subjects at baseline, and after RV-A16 infection, measured over time (c) Comparison of baseline and peak (i.e. the maximal concentration of MUC5B detected during the infection for each individual) levels of MUC5B and between subjects with COPD and healthy subjects. (d) Correlation of peak MUC5B with peak concentrations of inflammatory cells and cytokines in sputum. (e) Correlation between peak sputum MUC5B and peak virus loads. (f) Correlation of peak MUC5B with peak change from baseline levels of the indicated anti-microbial peptides in sputum. (g) Correlation of peak sputum MUC5B with bacterial loads assessed by 16S quantitative PCR. (h) Correlation of peak MUC5B with peak respiratory tract symptom scores. (i) Correlation of peak sputum MUC5B with peak expiratory flow rate (PEF) change from baseline during exacerbation. In (c) individual datapoints shown, analysed by Wilcoxon matched-pairs signed rank test for baseline vs peak and Mann Witney U test for comparison of peak values between COPD and healthy subjects. \*\**P*<0.01. In (d) - (i), correlation analysis used was nonparametric (Spearman’s correlation) performed on healthy volunteers and individuals with COPD pooled into a single group. In (g), 16S qPCR not measured in all patients due to lack of sample availability so data only shown where measured.

**Supplementary Figure 2:**
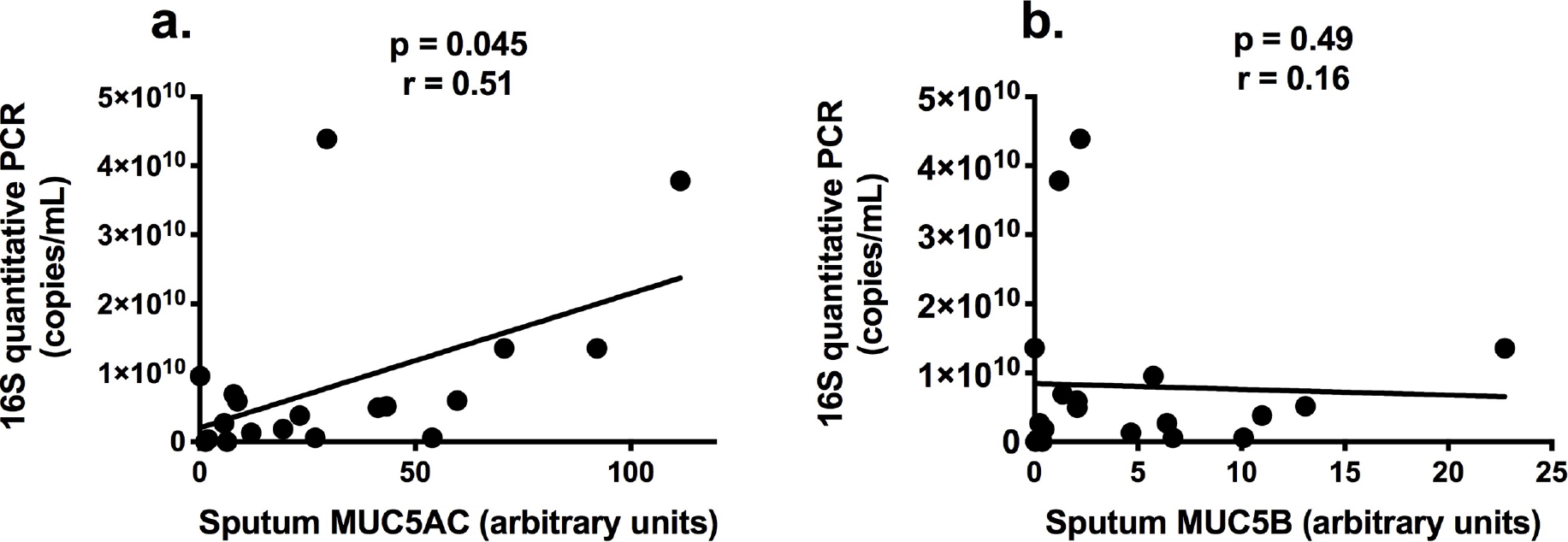
Correlation of sputum mucins with bacterial loads during naturally occurring virus-induced COPD exacerbations. (a) sputum Muc5ac and (b) sputum MUC5B correlation with 16S qPCR copies measured at 2 weeks following presentation with virus positive exacerbation in patients with COPD. Correlation analysis used was nonparametric (Spearman’s correlation).

**Supplementary Figure 3:**
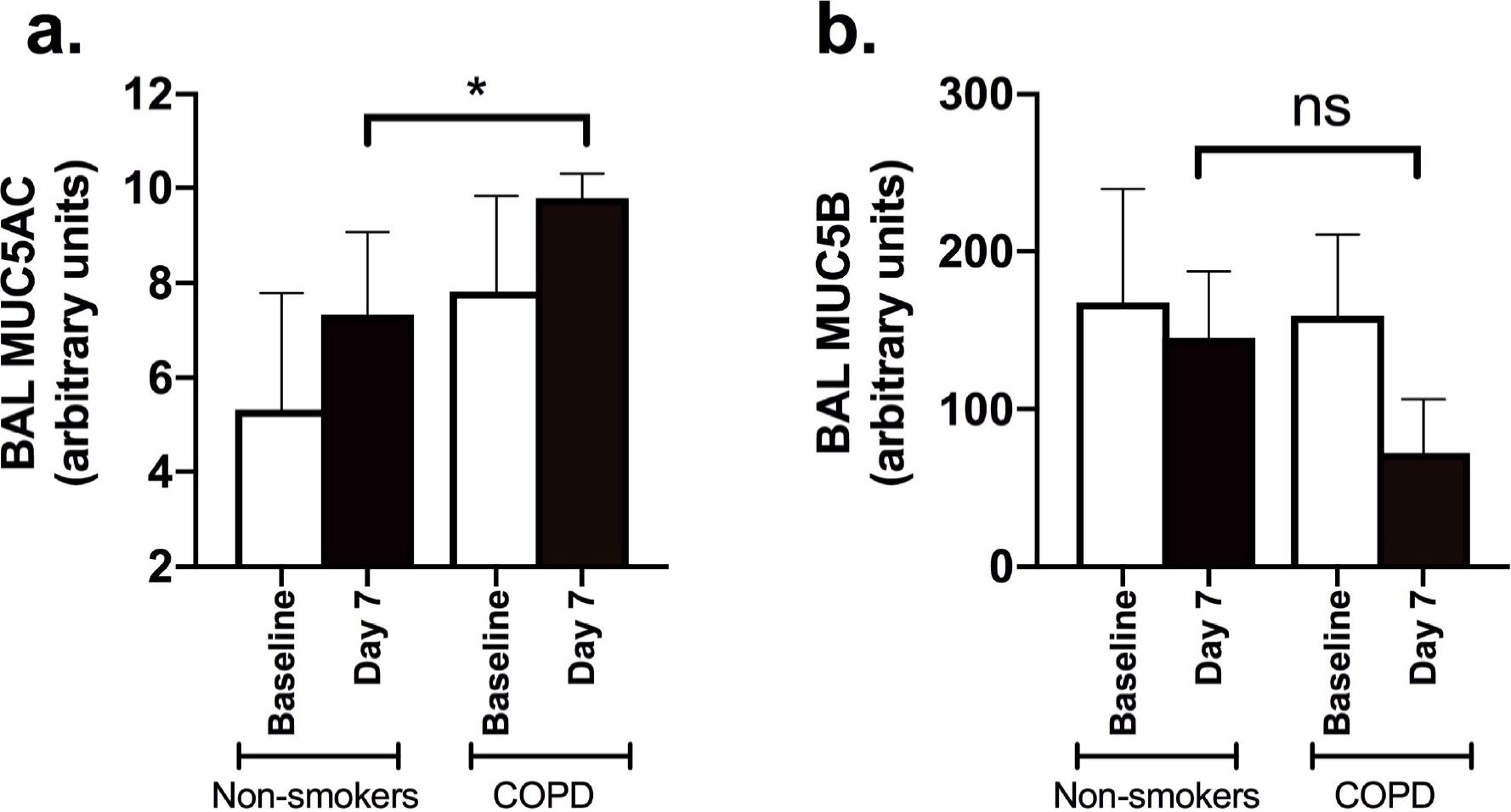
Sputum mucin concentrations in bronchoalveolar lavage during experimental rhinovirus infection in COPD and healthy subjects. (a) MUC5AC and (b) MUC5B concentrations in bronchoalveolar lavage samples at baseline and 7 days following experimental rhinovirus infection were measured by ELISA. Data expressed as median (+/− IQR) and analysed by Mann-WItney U test. *p<0.05. ns=non-significant.

**Supplementary Figure 4:**
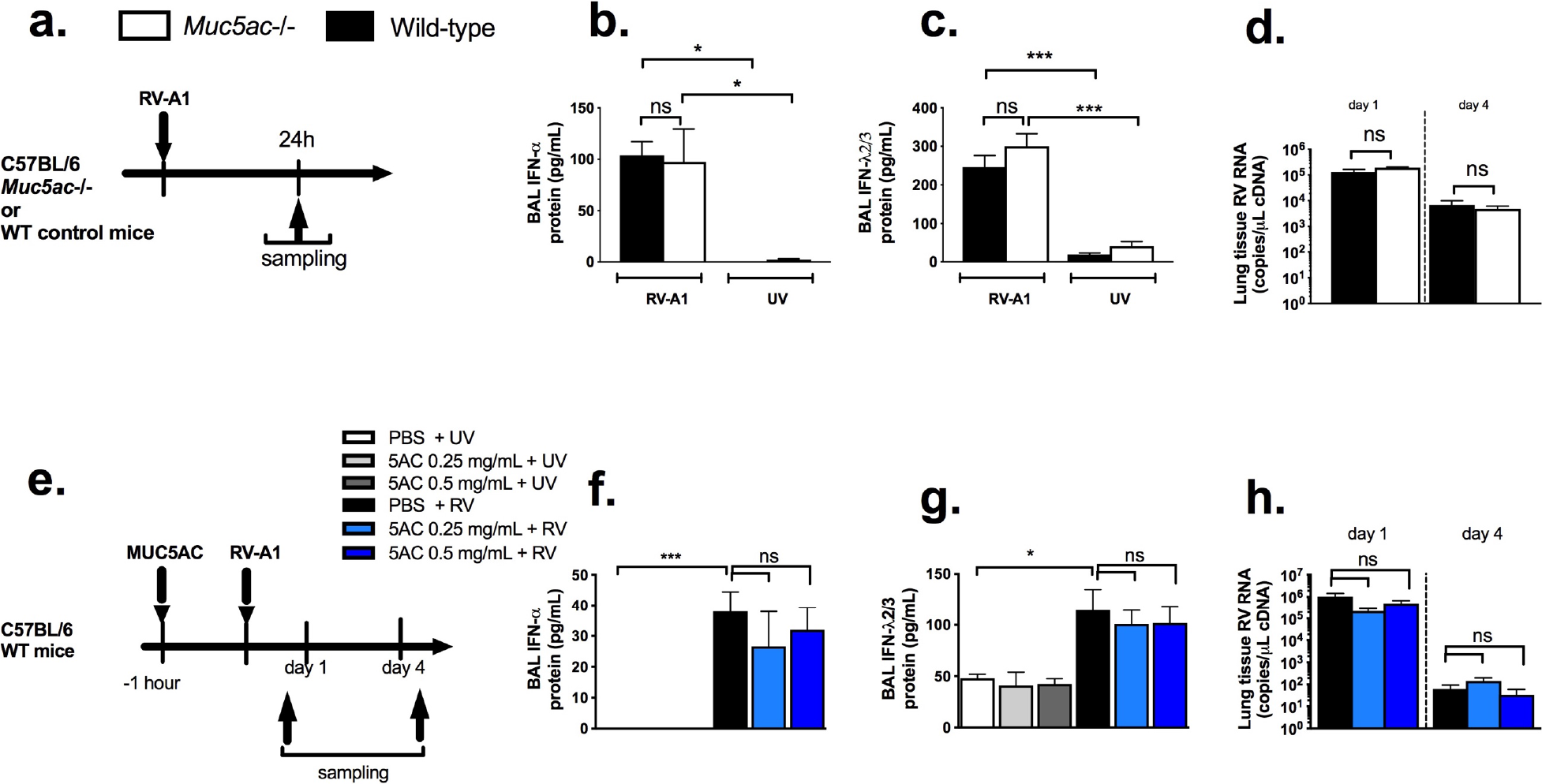
No effect of *Muc5ac* deletion or MUC5AC supplementation on anti-viral immunity or virus control. (a) Experimental outline. *Muc5ac*-/- or wild-type control C57BL/6 mice were intranasally infected with RV-A1 or UV inactivated RV-A1, (b) BAL concentrations of IFN-α and (c) IFN-λ2/3 proteins at day 1 post-infection. (d) Lung tissue rhinovirus RNA copies. (e) Experimental outline. C57BL/6 mice were treated intranasally with purified MUC5AC protein and additionally intranasally infected with RV-A1 or UV inactivated RV-A1. (f) BAL concentration of IFN-α and (g) IFN-λ2/3 proteins at day 1 post-infection. (h) Lung tissue rhinovirus RNA copies. All data expressed as mean (+/−S.E.M). Data analysed by one or two-way ANOVA with Bonferroni’s post-test. ns = non-significant *p<0.05; ***p<0.001.

**Supplementary Figure 5:**
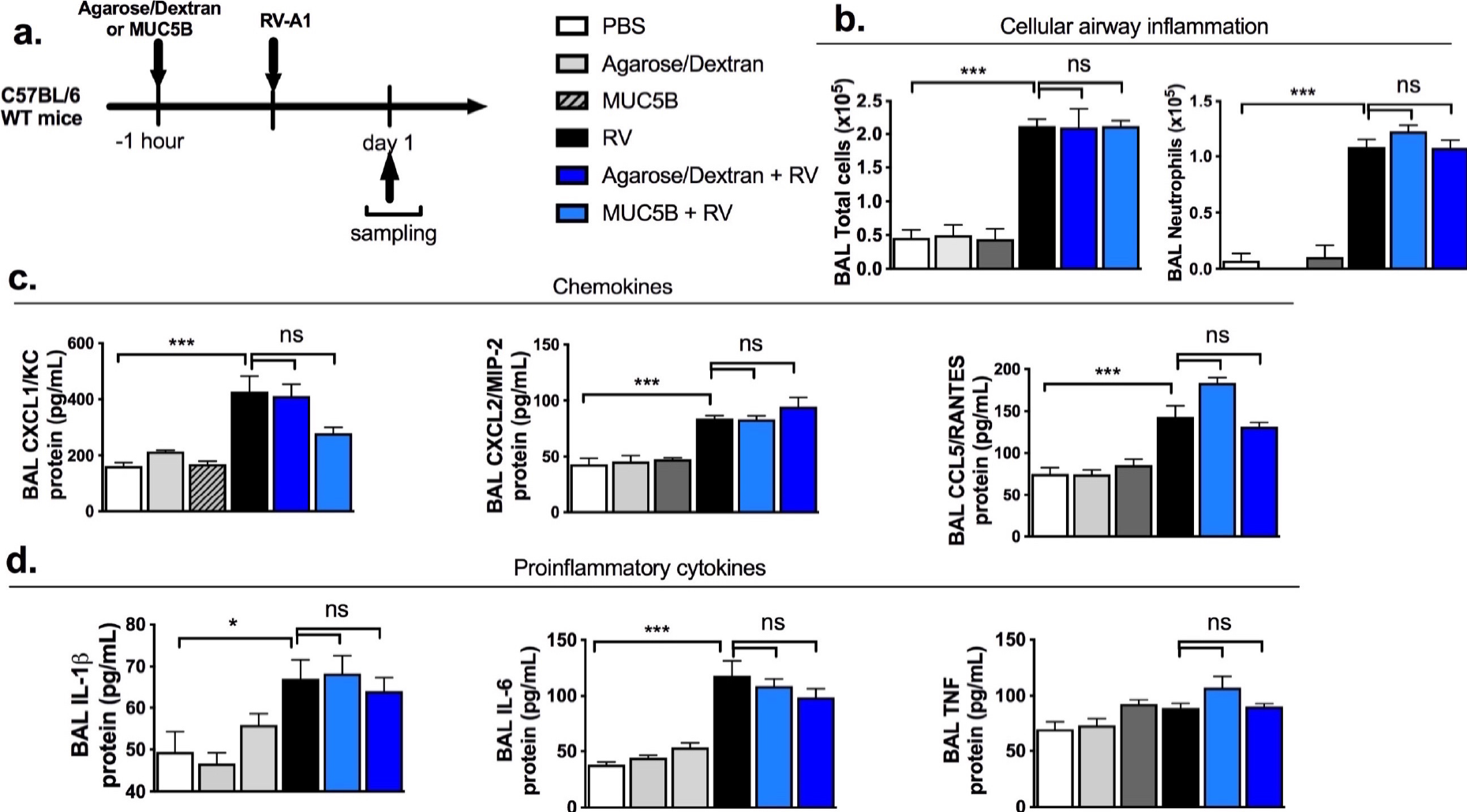
No effect of exogenous MUC5B or control polymer of agarose/dextran solution on rhinovirus-induced airway inflammation. (a) Experimental outline. C57BL/6 mice were treated intranasally with purified MUC5B protein or control polymer solution of agarose/dextran and additionally intranasally infected with RV-A1 or UV inactivated RV-A1. (b) BAL total cells and neutrophils at day 1 post-infection. (c) BAL concentrations of the indicated chemokines at day 1 post-infection. (d) BAL concentration of the indicated pro-inflammatory cytokines at day 1 post-infection. All data expressed as mean (+/−S.E.M). Data analysed by one-way ANOVA with Bonferroni’s post-test. \**P*<0.05; \*\*\**P*<0.001. ns = non-significant.

**Supplementary Figure 6:**
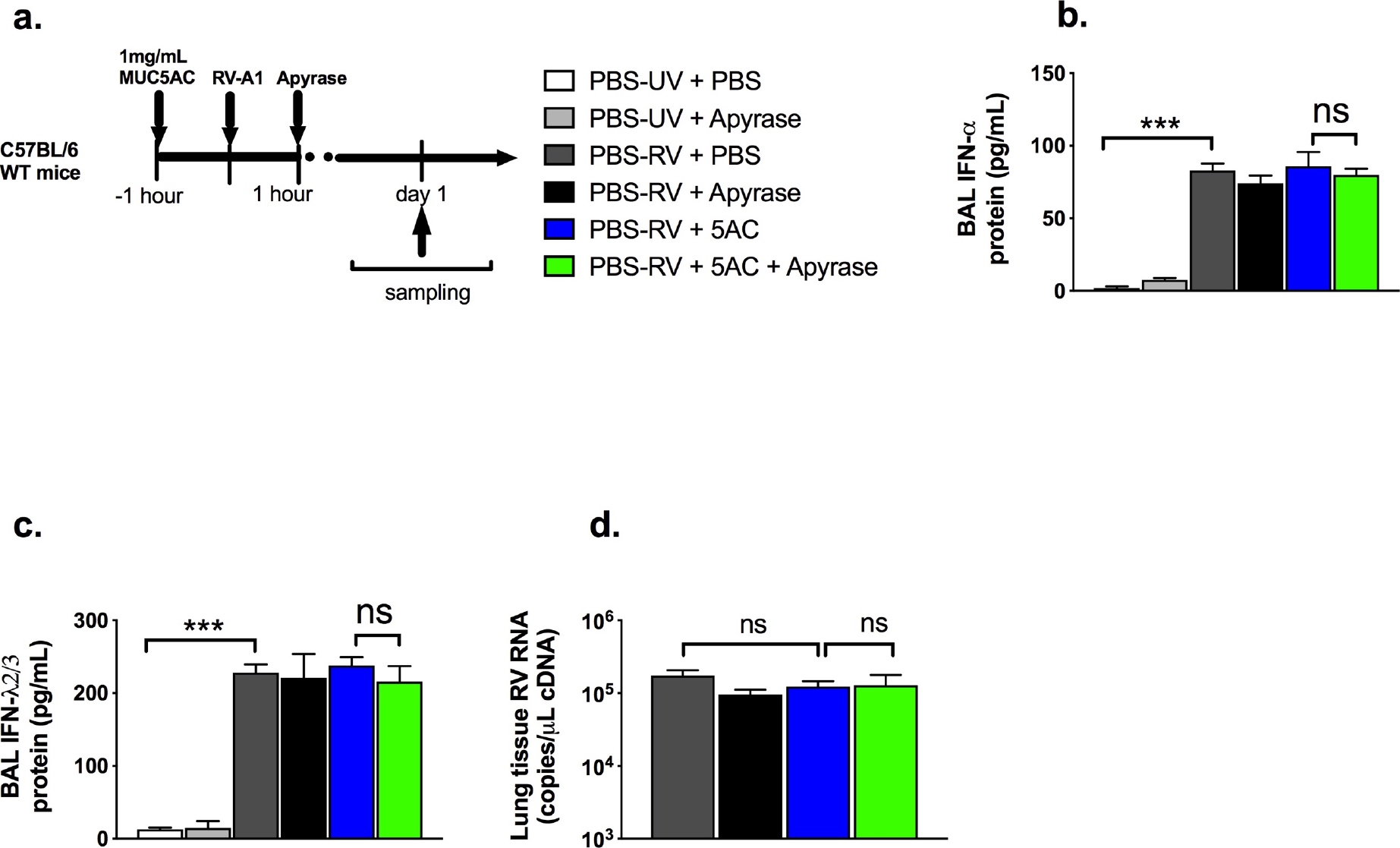
No effect of neutralization of airway ATP levels on anti-viral immunity or virus control. (a) Experimental outline. C57BL/6 mice were treated intranasally with purified MUC5AC protein or PBS control, infected with RV-A1 or UV inactivated RV-A1 and additionally treated with intranasal apyrase or vehicle control., (b) BAL concentrations of IFN-α and (c) IFN-λ2/3 proteins at day 1 post-infection. (d) Lung tissue rhinovirus RNA copies at dat 1 post-infection. All data expressed as mean (+/−S.E.M). Data analysed by one or two-way ANOVA with Bonferroni’s post-test. ns = non-significant *p<0.05; ***p<0.001.

**Supplementary Figure 7:**
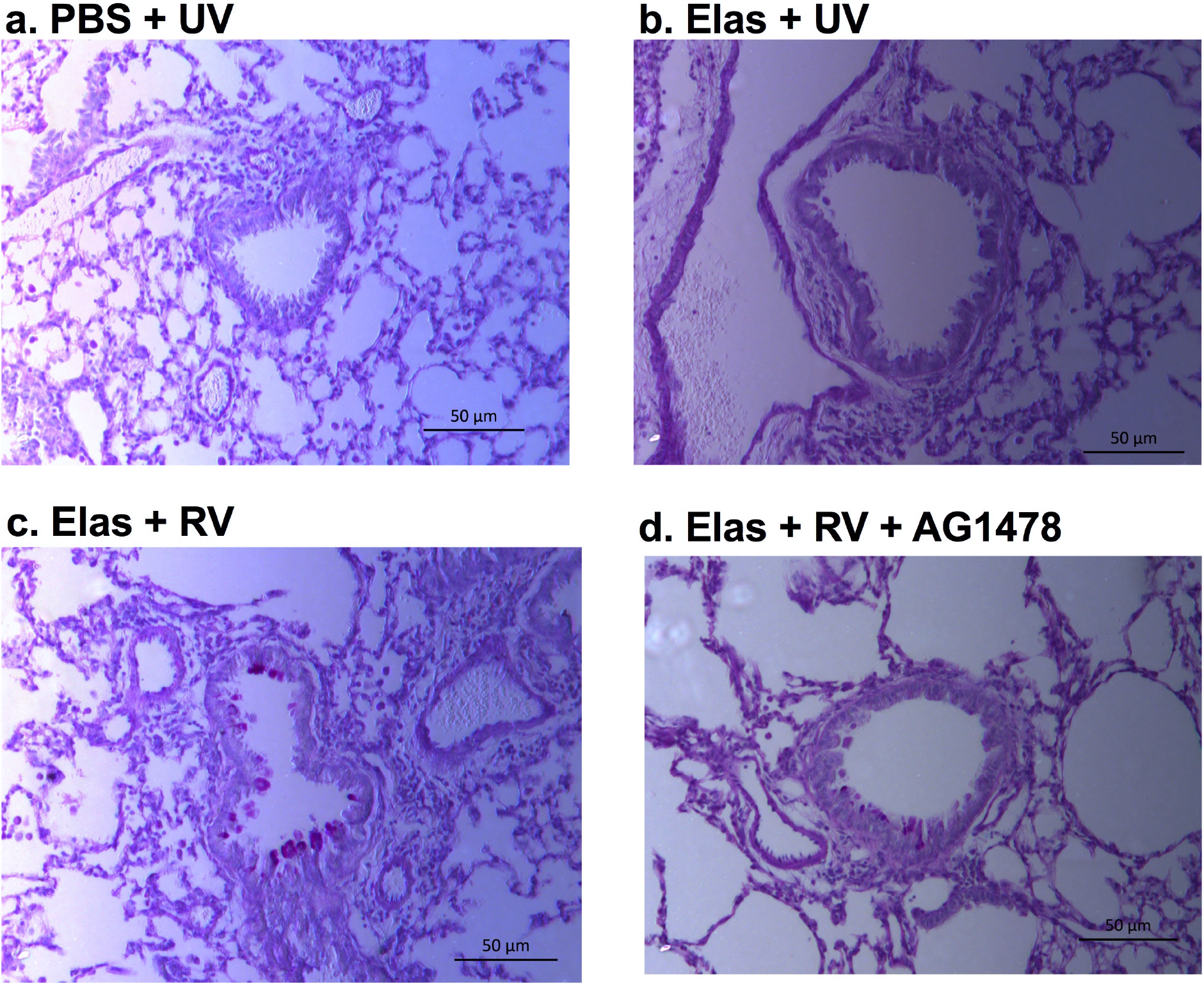
Period acid Schiff (PAS) stained lung sections in elastase treated mice infected with RV and treated with EGFR inhibitor AG1478 and control groups. C57BL/6 mice were treated intranasally with elastase or PBS control and additionally treated intraperitoneally with 50mg/kg of EGFR inhibitor AG1478, prior to challenge with rhinovirus (RV)-A1 or UV-inactivated RV-A1 (UV). At day 4 after RV challenge, lungs were formalin-fixed, paraffin-embedded and stained with periodic acid-schiff (PAS). Representative images shown from mice treated with (a) PBS + UV + vehicle (b) elastase + UV + vehicle (c) elastase + RV + vehicle (d) elastase + RV + AG1478.

**Supplementary Figure 8:**
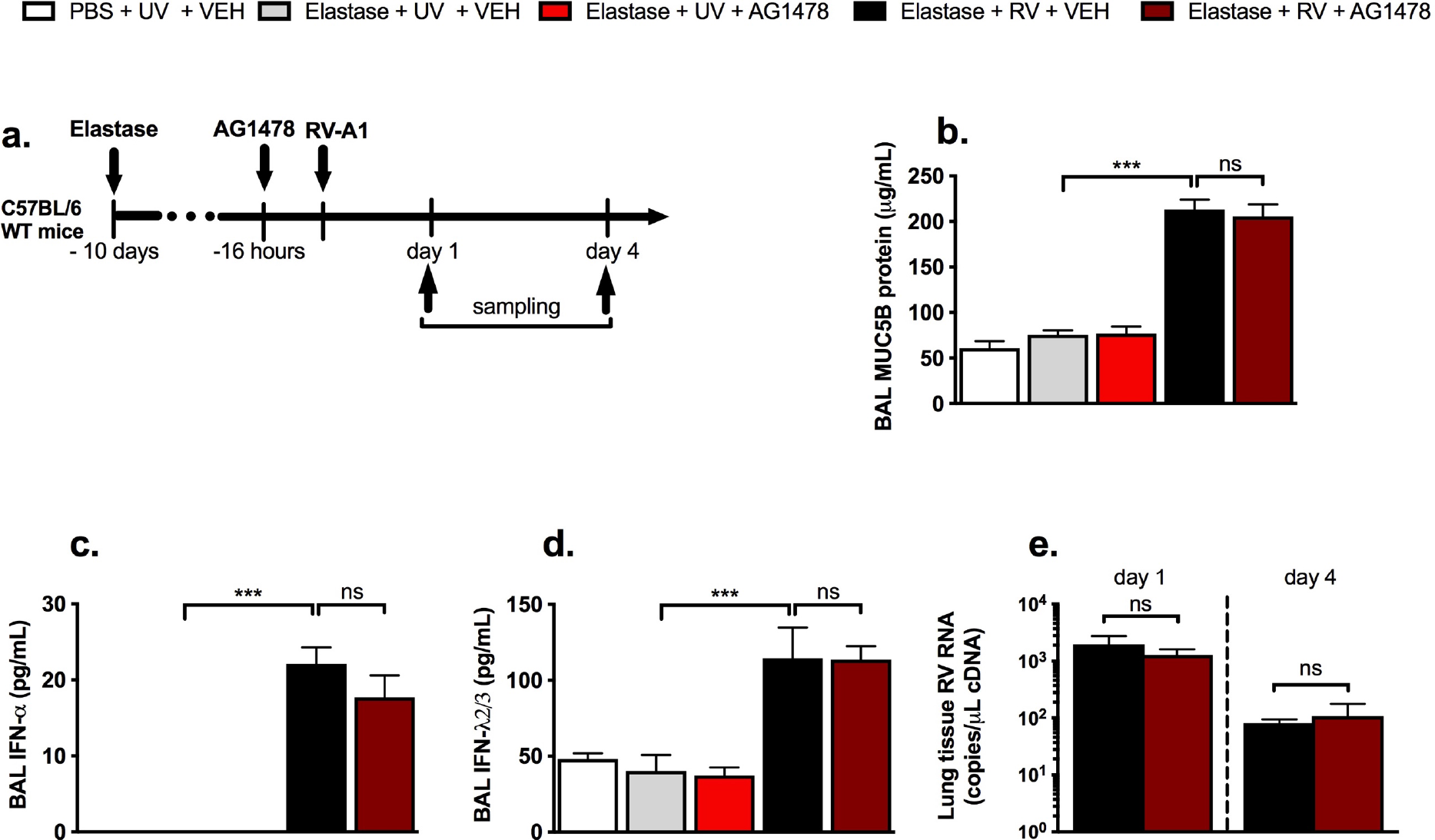
Effect of the EGFR inhibitor AG1478 on MUC5B, anti-viral immunity and virus control in a mouse model of rhinovirus-induced COPD exacerbation. (a) Experimental outline. C57BL/6 mice were treated intranasally with elastase or PBS control and additionally treated intraperitoneally with 50mg/kg of EGFR inhibitor AG1478, prior to challenge with rhinovirus (RV)-A1 or UV-inactivated RV-A1 (UV). (b) BAL MUC5B (c) IFN-α and (d) IFN-λ2/3 proteins at day 1 post-infection. (e) Lung tissue rhinovirus RNA copies. All data expressed as mean (+/−S.E.M). Data analysed by one or two-way ANOVA. ns = non-significant. ***p<0.001.

## Supplementary methods

### Purification of porcine gastric mucin MUC5AC

Previous studies revealed that the properties of commercially available mucins differ from key properties of native mucin systems as they, e.g. the lack of the ability to reduce friction^1-^3 or lead to cytotoxic effects^4^. These differences are attributed to the harsh conditions during the commercial purification process^5^.

As a consequence, here, the purification of porcine gastric MUC5AC is performed manually from fresh pig stomachs. An optimization of the original purification protocol from Celli *et.al*^6^ was described in detail previously^3^. In brief, crude mucus was obtained by gently scraping pig stomachs after rinsing them with tap water. The mucus was diluted 5-fold in 10 mM sodium phosphate buffer (pH 7.0, supplemented with 170 mM NaCl) and homogenized by stirring at 4 °C overnight. Cellular debris was removed via several centrifugation steps (8.300 × g for 30 min at 4 °C and 15.000 × g for 45 min at 4 °C) and a final ultracentrifugation step (150.000 × g for 1 h at 4 °C). Afterwards, the supernatant was separated chromatographically by size-exclusion chromatography (ÄKTA purifier system, GE Healthcare, equipped with a XK50/100 column packed with Sepharose 6FF), and the fractions containing the mucin glycoproteins were identified via PAS (periodic acid/Schiff’s base) staining and pooled. After a subsequent dialysis step against ultra-pure water via cross-flow filtration, the mucin-solution was concentrated, lyophilized and stored at −80 °C.

### Repurification of commercial bovine salivary mucin MUC5B

Lyophilized bovine salivary mucin MUC5B powder was purchased from Merck (499643, Darmstadt, Germany) and dissolved in 10 mM sodium phosphate buffer (pH 7.0, supplemented with 1 M NaCl) at a concentration of 1 mg mL^−1^ and the solution was rotated on a rotating incubator for 1 h at 4 °C. Further purification of salivary mucin MUC5B was conducted according to the protocol used for purifying gastric mucin MUC5AC as described above.

### Enzymatic removal of mucin-associated DNA

Since the gastric mucosa also contains DNA^7^, enzymatic removal of potential mucin-associated DNA was performed to prevent any DNA-related inflammation reactions. All following steps were carried out under a sterile hood. Lyophilized mucin purified from porcine gastric mucosa was treated with UV-light on ice for 1 hour. Subsequently, it was dissolved in 50 mM Tris-HCl buffer (sterile filtered, pH 7.5, supplemented with 10 mM MgCl_2_) at a concentration of 1 mg/mL and kept on ice. DNAse I from bovine pancreas (D5025, Sigma Aldrich, St. Louis, MO, USA) was dissolved in the same buffer at a concentration of 1 mg/mL. 50 μL of the DNAse solution were added per milligram of dissolved mucin. Afterwards, the mucin/DNAse solution was incubated at 37 °C for 18 hours in a tube shaker at 200 rpm. The treated mucin was then chromatographically separated from the enzyme and residual DNA fragments via size exclusion chromatography, subsequently desalted by means of cross-flow filtration and lyophilized (see *Purification of porcine gastric mucin* section above). The DNAse treated mucin was stored in lyophilized form at −80 °C.

### Visualization of mucin-associated DNA

Successful removal of DNA from MUC5AC was verified by electrophoretic separation of the mucin samples and subsequent visualization of associated DNA molecules via the DNA-binding dye SYBR Green I (S9430, Sigma Aldrich, St. Louis, MO, USA). Therefore, 40 μg of the mucin samples were mixed with Laemmli buffer, loaded on a polyacrylamide gel (Mini-Protean TGX Gels 4-20%, #45-1093, BioRad, Hercules, CA, USA) and separated (60 V for 10 min and 120 V until the dye front reached the end of the gel). The gel was stained using SYBR Green I afterwards (diluted 1:10.000 in TBE buffer) at room temperature under gentle shaking for 30 min. The DNA stain was visualized with a Gel Doc EZ Gel Documentation System (Biorad) with epi-illumination at a wavelength of 254 nm (**Fig. 1**). We additionally confirmed that DNA was undetectable in samples of MUC5AC protein using Quant-iT PicoGreen dsDNA reagent (Invitrogen, Carlsbad, CA), according to the manufacturer’s protocol, as previously described^8^.

**Figure 1:**
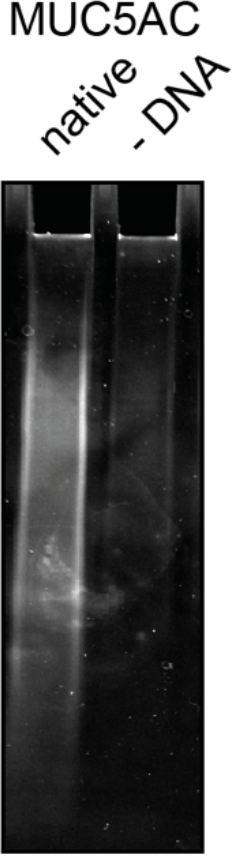
SDS-PAGE of native and DNAse-treated porcine gastric MUC5AC. DNA was removed from purified porcine gastric mucins by enzymatic treatment with DNAse I from bovine pancreas. Mucin samples were separated via gel-electrophoresis and subsequently stained with a SYBR Green nucleic acid gel stain to visualize associated DNA molecules (indicated by a white signal) Lane on right shows MUC5AC protein with DNA removed.

